# Comprehensive Genome Based Analysis of *Vibrio parahaemolyticus* for Identifying Novel Drug and Vaccine Molecules: Subtractive Proteomics and Vaccinomics Approach

**DOI:** 10.1101/2020.04.17.045849

**Authors:** Mahmudul Hasan, Kazi Faizul Azim, Abdus Shukur Imran, Ishtiak Malique Chowdhury, Shah Rucksana Akhter Urme, Md Sorwer Alam Parvez, Md Bashir Uddin, Syed Sayeem Uddin Ahmed

## Abstract

Multidrug-resistant *Vibrio parahaemolyticus* has become a significant threat to human health as well as aquaculture, prioritizing the development of effective drug and vaccine candidates. Hence, the study was designed to identify novel therapeutics using a comprehensive genome-based analysis of *V. parahaemolyticus.* From *V. parahaemolyticus* proteome, a total of 4822 proteins were investigated in order to find out effective drug and vaccine targets. A range of diverse subtractive proteomics approaches – namely, identification of human non-homologous and pathogen-specific essential proteins, druggability and ‘anti-target’ analysis, prediction of subcellular localization, human microbiome non-homology screening, analysis of virulence factors, protein-protein interactions studies. Among 16 novel cytoplasmic proteins, ‘VIBPA Type II secretion system protein L’ and ‘VIBPA Putative fimbrial protein Z’ were allowed to molecular docking with 350 human metabolites, which revealed that Eliglustat, Simvastatin and Hydroxocobalamin were the top drug molecules considering free binding energy. On the contrary, ‘Sensor histidine protein kinase UhpB’ and ‘Flagellar hook-associated protein of 25 novel membrane proteins were subjected to T and B cell epitope prediction, antigenicity testing, transmembrane topology screening, allergenicity and toxicity assessment, population coverage analysis and molecular docking were adopted to generate the most immunogenic epitopes. Three subunit vaccines were constructed by the combination of highly antigenic epitopes along with suitable adjuvant, PADRE sequence and linkers. The designed vaccine constructs (V1, V2, V3) were analyzed by their physiochemical properties and molecular docking with MHC molecules that suggested the superiority of construct V1. Besides, the binding affinity of human TLR1/2 heterodimer and construct V1 was also biologically significant. The vaccine-receptor complex exhibited deformability at a minimum level that also strengthened our prediction. The optimized codons of the designed construct was cloned into pET28a(+) vector of *E. coli* strain K12. However, the predicted drug molecules and vaccine constructs could be further studied to combat *V. parahaemolyticus* associated infections.

## Introduction

*Vibrio parahaemolyticus*, a highly reported pathogenic bacteria of aquatic environment, belongs to vibrionaceae family has emerged as the leading cause of seafood-associated gastroenteritis and a significant hazard for global aquaculture (Tang et al., 2014, Ghenem et al., 2017, Chen et al., 2017, Letchumanan et al., 2014). The overgrowing population, with increased purchasing power worldwide, has enhanced the demand for and export potential of seafood, resulting in the steady expansion of the aquaculture industry (Rico et al., 2012). However, the sector has continuously been challenged by aquatic animal health problems, which are a significant constraint to the development of this sector (Bondad-Reantaso et al., 2005). Besides, Antimicrobial resistance (MDR) has been recognized as an essential global threat issue to food safety (Food and Agriculture Organization, 2016). The continuous and inappropriate use of antibiotics in the aquaculture industry favors the development of a variety of resistant isolates and the dissemination of resistance genes within the bacterial population (Tendencia and de laPena, 2001). *V. parahaemolyticus* has been reported to show multidrug resistance during aquaculture production (Vaseeharan et al., 2005; Han et al., 2007, Yang et al., 2017), which raised the concern about public health and economic threat of this bacterium (Lesley et al., 2011; Noorlis et al., 2011; Osunla et al., 2017).

Though *V. parahaemolyticus* was first isolated in 1952, reports demonstrated the recent outbreaks of *V. parahaemolyticus* are more severe (Jung, 2018; Liu et al., 2015; Jang et al., 2013). On the recent outbreak in the city of Osaka (Japan), acute gastroenteritis was reported in 272 individuals, 20 of whom died (Daniels et al., 2000). To date, *V. parahaemolyticus* has been responsible for 20– 30% of food poisoning cases in Japan and sea foodborne diseases in many Asian countries (Alam et al., 2002). A total 802 outbreaks of food-borne diseases have been reported in 13 of the coastal provinces of eastern China, causing more than 17,000 individuals to become ill (Wang et al., 2011a; Wu et al., 2014), where *V. parahaemolyticus* attributed the most significant number (40.1%) of these cases (Liu et al., 2006, Nair et al., 2007, Chao et al., 2011). The leading cause of human gastroenteritis associated with seafood consumption in the United States is *V. parahaemolyticus* (Kaysner and De Paola, 2001; Newton et al., 2012). Centers for Disease Control and Prevention (CDC) declared it as a significant foodborne bacterium compared to other *Vibrio* species, which was responsible for approximately 34,664 foodborne cases annually in the USA (Scallan et al., 2011; Huang et al., 2016).

The food poisoning caused by *V. parahaemolyticus* usually occurs in summer and is predominantly associated with different kinds of seafood, including crab, shrimp, shellfish, lobster, fish and oysters (Cruz et al, 2015; Letchumanan et al., 2015). *V. parahaemolyticus* is usually found in a free-swimming state, with its motility conferred by a single polar flagellum affixed to inert and animate surfaces including zooplankton, fish, shellfish or any suspended matter underwater (Gode-Potratzetal, 2011). Among the whole range of seafood, shellfish is regarded as a high-risk food because it is infested with large populations of bacteria, including *V. parahaemolyticus* (Peng et al., 2010; China Statistical Yearbook, 2012). Illness is inevitable, once consumers eat undercooked contaminated seafood (Rahimi et al., 2010). The symptoms of the disease include diarrhea, vomiting, abdominal pain, nausea and low-grade fever (Ham and Orth, 2012). In most cases, the disease is self-resolving. However, *V. parahaemolyticus* may cause a more debilitating and dysenteric form of gastroenteritis (Levin, 2006). Uncommonly, in immunocompromised patients, it may progress into a life-threatening fulminant necrotizing fasciitis characterized by rapid necrosis of subcutaneous tissue (Ahmad et al., 2013). In rare cases, *V. parahaemolyticus* causes septicemia, which is also associated with a high mortality rate (Zhang and Orth, 2013). Also, *V. parahaemolyticus* is one of the major pathogens of cultured mud crabs and cause acute hepatopancreatic necrosis disease (AHPND) in shrimp (Xiao et al., 2017). Usually, 99% of clinical *V. parahaemolyticus* isolates are known to be pathogenic, whereas the majority of the environmental isolates are non-pathogenic (Yu et al., 2013; Sudha et al., 2014). Nonetheless, around 0–6% of the environmental isolates are identified as pathogenic carrying virulence genes (Letchumanan et al., 2014). During infection, *V. parahaemolyticus* uses the adhesion factors to bind to the fibronectin and phosphatidic acid on the host cell, thus releasing different effectors and toxins into the cytoplasm, causing cytotoxicity and serious diseases (Gode-Potratz et al., 2011).

Many antibiotics are no longer effective in hospitals to treat *V. parahaemolyticus* infections (Tan et al., 2016, Jun et al., 2014, Lin et al., 2017). First-generation antibiotics, including ampicillin are extensively used in aquaculture resulting in reduced susceptibility and low efficacy of ampicillin for *Vibrio* sp. treatment (Sudha et al., 2014, Han et al., 2007). Literature also reported higher resistance to third-generation antibiotics such as cephalosporin, cefotaxime, carbapenems and ceftazidime by *V. parahaemolyticus* isolates (Lee et al., 2018; Jun et al., 2012; Letchumanan et al., 2015) which enhanced the necessity of searching safe and more effective drugs for combating infections caused by *V. parahaemolyticus* in the future. However, the development of new antibiotics is difficult and time-consuming. Recent progress in the field of computational biology and bioinformatics has generated various in silico analysis and drug designing approaches. Thus eliminating the time and cost involved in the early trial phase before going into the drug development phase (Barh et al., 2011). Subtractive genomics is one such in silico strategy that helps to facilitate the selection, processing, and development of strain-specific drugs against various pathogens (Azim et al., 2019). It can be utilized to identify drug targets based on the determination of essential and nonhomologous proteins within the pathogenic organism (Barh and Kumar, 2009; Hosen et al., 2014). Various novel drug targets have already been successfully identified for *S. typhi meningitides*sero group B (Perumalet al., 2007; Barh and Kumar, 2009) using the mentioned approach.

Moreover, *in silico* docking studies between the identified drug targets and existing drugs with slight modification may lead to the discovery of novel drugs for the treatment of infections (Pagadala et al., 2017; Wong, 2015; Ferreira et al., 2015; Yuriev et al., 2013). As a result, a wide range of drug targets and lead compounds can be identified before laboratory experimentation, to save time and money. The study was designed to employ a comprehensive genome-based analysis of *Vibrio parahaemolyticus* for identifying novel therapeutic targets as well as suitable drug and vaccine molecules through subtractive proteomics and vaccinomics approaches.

## Results

Various bioinformatics tools and databases were used to analyze the entire proteome of *V. parahemolyticus* through subtractive genomics and vaccinomics approach. The step by step results (or workflow in Fig. 1 and Fig. 2) from the complete computational analysis was presented in Table 1.

**Table 1:** Subtractive genomic analysis scheme

| Sl. No. | Steps | No. of proteins | Protein sets |
| --- | --- | --- | --- |
| 1. | Total number of proteins | 4822 | Set 0 (Supplementary file 1) |
| 2. | Non-paralogous (>60% identical) in CD-Hit | 4729 | Set 1 (Supplementary file 2) |
| 3. | proteins with >100 amino acids | 4123 | Set 2 (Supplementary file 3) |
| 4. | Number of proteins nonhomologous to *H. sapiens* using BLASTp (E value $10^{-3}$) | 3164 | Set 3 (Supplementary file 4) |
| 5. | Essential proteins in DEG 15.2 server (E value $\leq 10^{-100}$, Bit score >100) | 1107 | Set 4 (Supplementary file 5) |
| 6. | Essential Proteins involved only in unique metabolic pathways (KAAS at KEGG) | 96 | Set 5 (Supplementary file 9) |
| 7. | Essential proteins found to be novel in Drug Bank 5.1.0 (using default parameters) | 41 | Set 6 (Supplementary file 10) |
| 8. | Novel drug target proteins non-homologous to ‘anti-targets’ using BLASTp (E value <0.005, Identity <25%, Query length >30%) | 41 | Set 7 (Supplementary file 10) |
| 9. | Essential cytoplasmic proteins using PSORTb, CELLO, ngLOC, PSLpred | 16 | Set 8 (Supplementary file 12) |
| 10. | Proteins showing <45% similarity with human microflora proteins | 9 | Set 9 (Supplementary file 13) |
| 11. | Identified virulence associated novel proteins by VFDB analysis | 2 | Set 10 (Table 4) |
| 12. | Essential membrane proteins using PSORTb, CELLO, ngLOC, PSLpred | 25 | Set 11 (Supplementary file 14) |
| 13. | Identified vaccine targets with less similarity with human microflora proteins and antigenicity | 2 | Set 12 (Table 7) |

**Figure 1.**
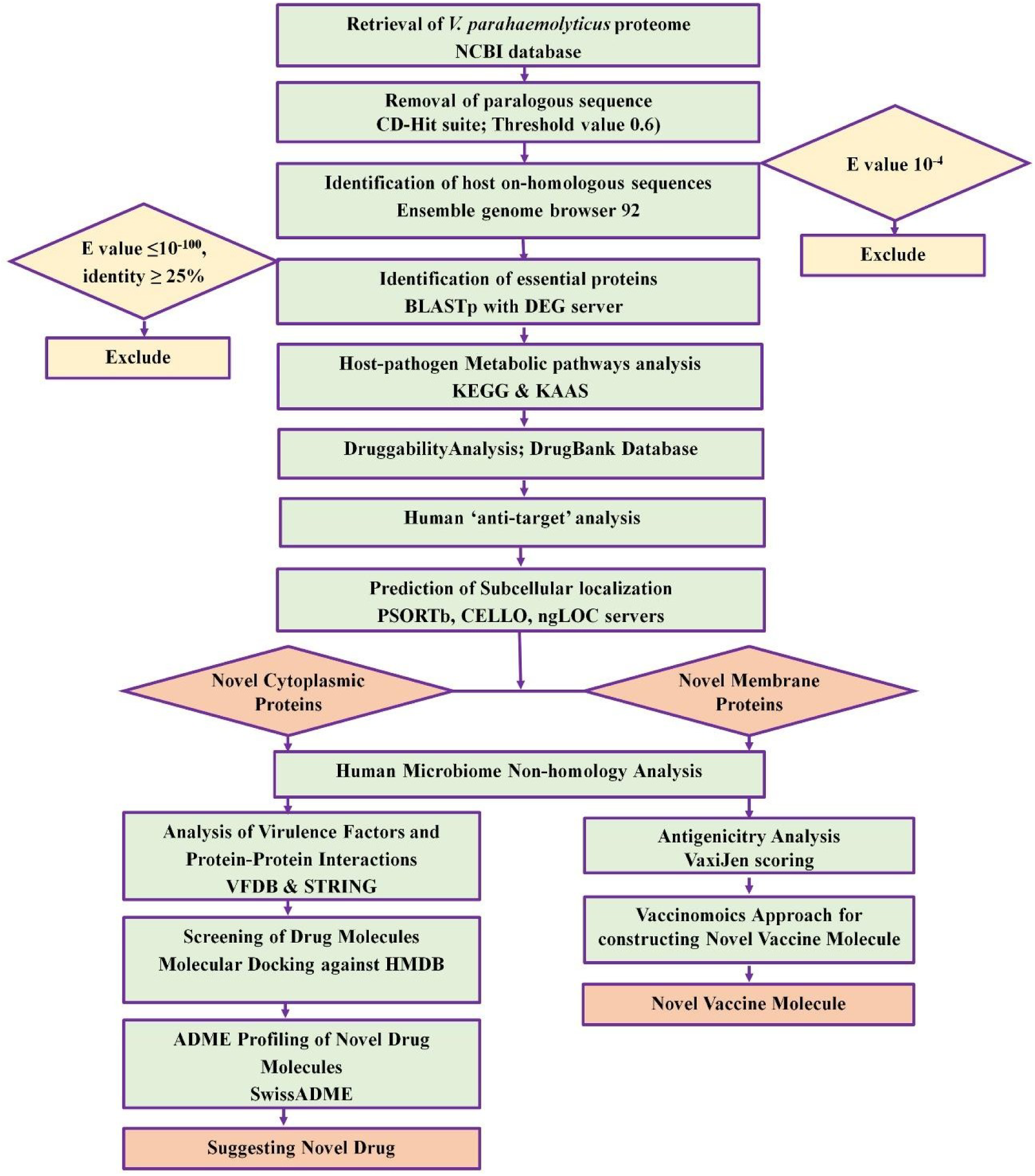
Proteome exploration of *Vibrio parahaemolyticus* to identify novel drug targets.

**Figure 2.**
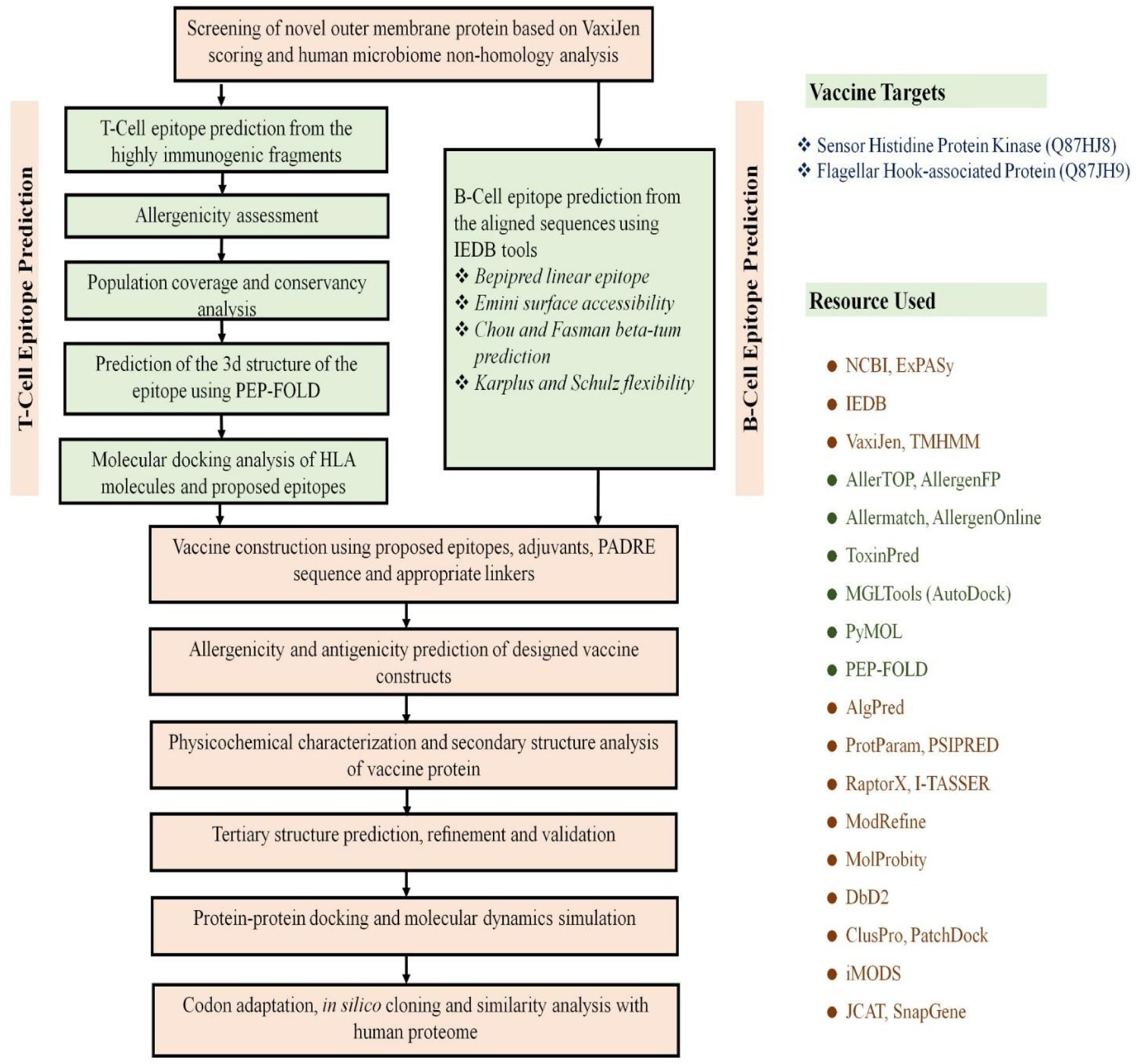
Flow chart summarizing the protocols over multi-epitope subunit vaccine development against *V. parahaemolyticus* through reverse vaccinology approach.

### Retrieval of complete proteome and identification of essential proteins

The whole proteome of *V. parahemolyticus* strain O3:K6 was extracted from NCBI database (Supplementary file 1) containing 4822 proteins (Set 0). Paralogous protein sequences of the pathogen. A total of 93 paralogous sequence above >60% similarity were identified through the CD-hit server and removed, leaving 4729 non-paralogous protein sequences in Set 1 (Supplementary file 2). Among these proteins, proteins with >100 residues (4123 proteins) (Set 2) were only considered (Supplementary file 3) for further analysis. Again, proteins showing significant similarity with human RefSeq proteins (1143 proteins) were excluded from the list designated as Set 3 (Supplementary file 4). Analysis of remaining proteins through the DEG server revealed only proteins that are essential (Set 4) for the survival of the pathogen (Supplementary file 5).

### Analysis of Metabolic pathways

The KEGG server contains 131 metabolic pathways for *V. parahaemolyticus* (Supplementary file 6) and 325 pathways for humans (Supplementary file 7). Through manual comparison, 40 metabolic pathways were found to be pathogen-specific and are provided in Supplementary file 8. Proteins involved in these unique pathways can be selected as drug targets. Non-homologous essential proteins subjected to BLASTp in the KAAS server at KEGG revealed that 96 proteins among 1107 assigned both KO (KEGG Orthology) and metabolic pathways that further deputed as Set 5 (Supplementary file 9).

### Druggability analysis and identification of novel drug targets

Only 56 proteins showed similarity with the available drug targets, while the remaining 41 showed no hit. These 41 proteins (Set 6) were considered as novel drug targets which include both cytoplasmic and membrane proteins (Supplementary file 10). Besides, the results of druggable proteins are provided in supplementary table 1. Furthermore, the other 41 proteins were considered as novel therapeutic targets and subjected to human ‘anti-targets’ analysis.

### ‘Anti-target’ analysis and prediction of subcellular localization

A total of 210 ‘anti-targets’ reported in the literature were fetched from NCBI (Supplementary file 11). All novel drug target proteins were successfully screened through BLASTp analysis, and no evidence of similarity was seen. Hence, all these novel drug target proteins were listed for human microbiome analysis considering non-homologous to host ‘anti-targets’ (Supplementary file 10). Moreover, The results of subcellular localization analysis by four servers are provided in Supplementary file 11. The result revealed that among 41 specific proteins involved in pathogen-specific pathways, 16 were cytoplasmic proteins assigned as Set 8 (Supplementary file 12, Table 2), while the remaining 25 sequences were membrane proteins.

**Table 2:**
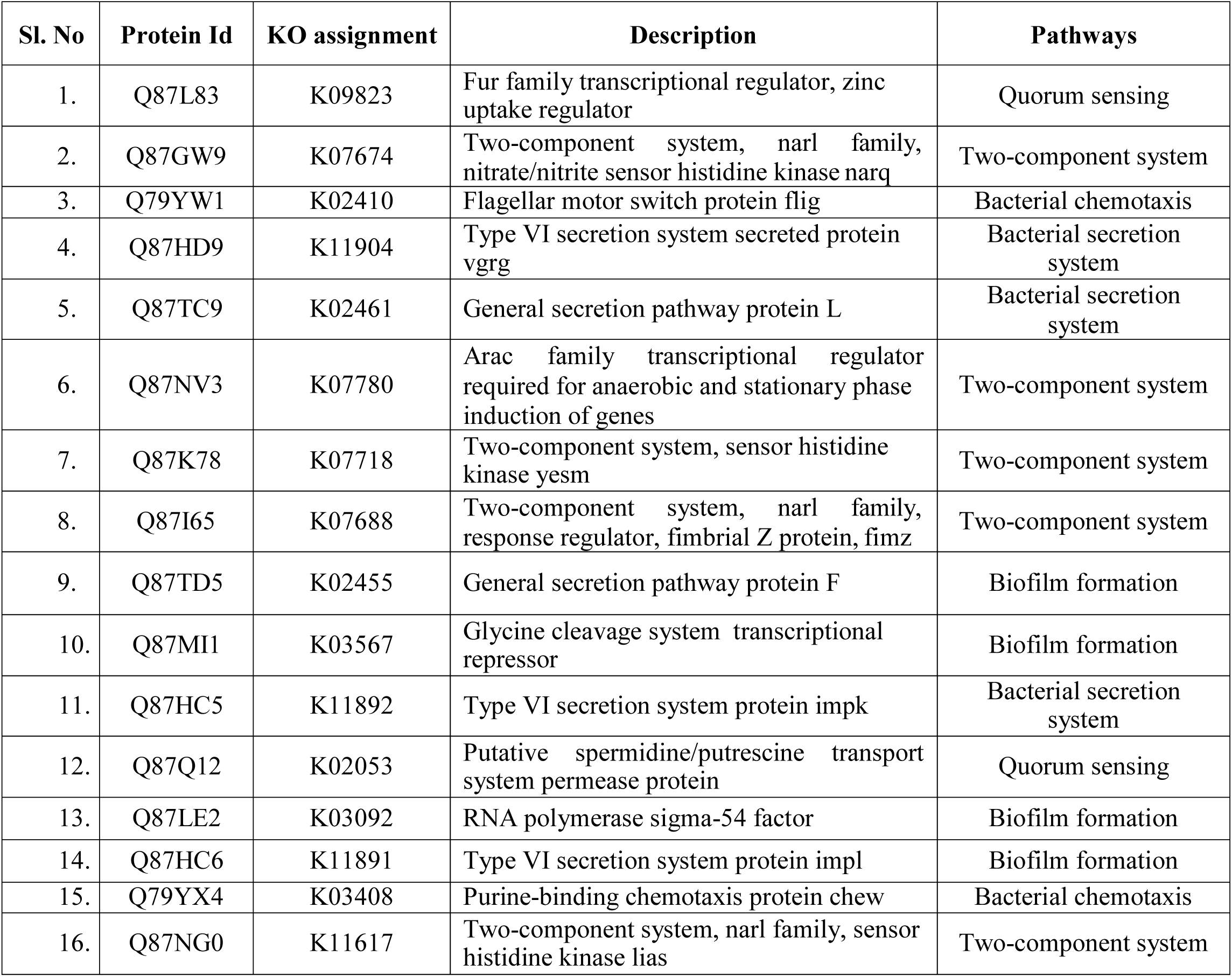
Pathogen specific essential cytoplasmic proteins as novel therapeutic targets.

### Human microbiome non-homology analysis

Cytoplasmic proteins showing <45% similarity with reported human microbiome proteins were selected for protein-protein interaction analysis, whereas membrane proteins were selected for further vaccine candidacy. However, microbiome analysis revealed that a total of 9 proteins (Set of the pathogen showed <45% similarity with human microflora (Supplementary Table 2, Supplementary file 13).

### Analysis of virulence factors (VF’s) and protein-protein interactions studies (PPIs)

From 9 cytoplasmic novel proteins, five uncharacterized proteins were removed and the remaining four proteins were considered for VFDV analysis. The VFDB result showed that two proteins (Set i.e., VIBPA Type II secretion system protein L (Q87TC9) and VIBPA Putative fimbrial protein Z (Q87I65) were associated with virulence of *V. parahaemolyticus* (Table 3). These proteins were subjected to protein-protein interaction study. STRING v10.5 revealed that Type II secretion system protein L confers interactions with nine proteins (Fig. 3A), while putative fimbrial protein Z exhibits interactions with three other proteins (Fig. 3B). These proteins are mainly responsible for protein transport, involved in biofilm formation and bacterial secretion system, or act as regulatory proteins (e.g., transcription regulator, signal transduction response regulator).

**Table 3:** Predicted therapeutic targets (novel cytoplasmic proteins) showing virulent properties.

| Novel drug targets | Accession Id | VFDB analysis | Interacted proteins |
| --- | --- | --- | --- |
| VIBPA Type II secretion system protein L | Q87TC9 | 5 hits | 9 |
| VIBPA Putative fimbrial protein Z | Q87I65 | 4 hits | 3 |

**Figure 3.**
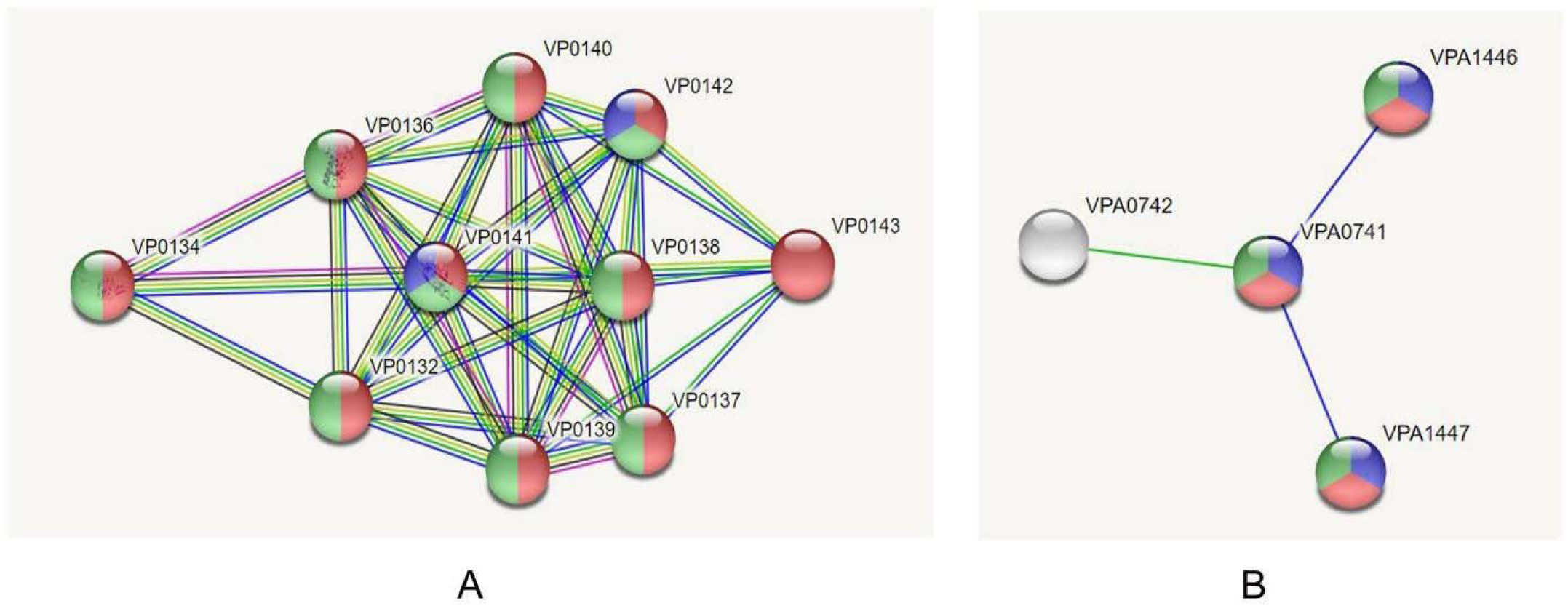
Investigation of Protein-Protein Interactions through STRING v10.5 server; (A) UDP-N-acetylmuramoyl-L-alanyl-D-glutamate--2,6-diaminopimelate ligase (murE), (B) Trigger factor (tig)

### Screening of drug molecules against novel cytoplasmic proteins

A total of 335 unique metabolites were retrieved from Human Metabolites Database for docking analysis against predicted therapeutic drug targets (novel cytoplasmic proteins). Docking scores were analyzed to screen the top drug candidates with the lowest binding energy (Supplementary Table 3). Among top 10 metabolites (Table 4), Eliglustat (DB09039) was found superior in terms of free binding energy for both protein targets, followed by Simvastatin (DB09039) and Hydroxocobalamin (DB00200) for VIBPA Type II secretion system protein L (Q87TC9) and VIBPA Putative fimbrial protein Z (Q87I65) respectively. ADME analysis was performed to get an insight into how the predicted pharmaceuticals will interact with the body as a whole (Table 5).

**Table 4:** Top 10 metabolites predicted as suitable drug candidates against VIBPA Type II secretion system protein L and VIBPA Putative fimbrial protein Z.

| Novel Targets | HMDB ID | Binding Energy | Name | Drug Bank ID | Drug Name | Drug Group |
| --- | --- | --- | --- | --- | --- | --- |
| *VIBPA Type II secretion system protein L (Q87TC9)* | HMDB0004971 | -10.6 | Glucosylceramide (d18:1/16:0) | DB09039 | Eliglustat | Approved |
|  | HMDB0004972 | -10.5 | Glucosylceramide (d18:1/18:0) | DB09039 | Eliglustat | Approved |
|  | HMDB0004970 | -10.3 | Glucosylceramide (d18:1/9Z-18:1) | DB09039 | Eliglustat | Approved |
|  | HMDB0004976 | -10.3 | Glucosylceramide (d18:1/26:1(17Z)) | DB09039 | Eliglustat | Approved |
|  | HMDB0008646 | -10.3 | PC(22:4(7Z,10Z,13Z,16Z)/22:4(7Z,10Z,13Z,16Z)) | DB00641 | Simvastatin | Approved |
|  | HMDB0011300 | -10.1 | PC(P-18:1(11Z)/P-18:1(11Z)) | DB00334 | Olanzipine | Approved, Investigational |
|  | HMDB0008443 | -9.7 | PC(20:4(5Z,8Z,11Z,14Z)/20:4(5Z,8Z,11Z,14Z)) | DB00641 | Simvastatin | Approved |
|  | HMDB0008138 | -9.6 | PC(18:2(9Z,12Z)/18:2(9Z,12Z)) | DB00641 | Simvastatin | Approved |
|  | HMDB0004973 | -9.5 | Glucosylceramide (d18:1/20:0) | DB09039 | Eliglustat | Approved |
|  | HMDB0010348 | -9.5 | Dehydroepiandrosterone 3-glucuronide | DB01708 | Prasterone | Approved, Investigational, Nutraceutical |
| *VIBPA Putative fimbrial protein Z (Q87I65)* | HMDB0004972 | -9.6 | Glucosylceramide (d18:1/18:0) | DB09039 | Eliglustat | Approved |
|  | HMDB0002308 | -9.5 | Hydroxocobalamin | DB00200 | Hydroxocobalamin | Approved |
|  | HMDB0004970 | -9.3 | Glucosylceramide (d18:1/9Z-18:1) | DB09039 | Eliglustat | Approved |
|  | HMDB0004971 | -9.3 | Glucosylceramide (d18:1/16:0) | DB09039 | Eliglustat | Approved |
|  | HMDB0002174 | -9.1 | Cobalamin | DB14098 | Cobalamin | Experimental |
|  | HMDB0008646 | -8.9 | PC(22:4(7Z,10Z,13Z,16Z)/22:4(7Z,10Z,13Z,16Z)) | DB00641 | Simvastatin | Approved |
|  | HMDB00060546 | -8.9 | Norbuprenorphine | DB01026 | Ketoconazole | Approved, Investigational |
|  | HMDB00041936 | -8.7 | Morphine-3-glucuronide | DB00295 | Morphine | Approved, Investigational |
|  | HMDB0004974 | -8.6 | Glucosylceramide (d18:1/22:0) | DB09039 | Eliglustat | Approved |
|  | HMDB0008443 | -8.5 | PC(20:4(5Z,8Z,11Z,14Z)/20:4(5Z,8Z,11Z,14Z)) | DB00641 | Simvastatin | Approved |

**Table 5:**
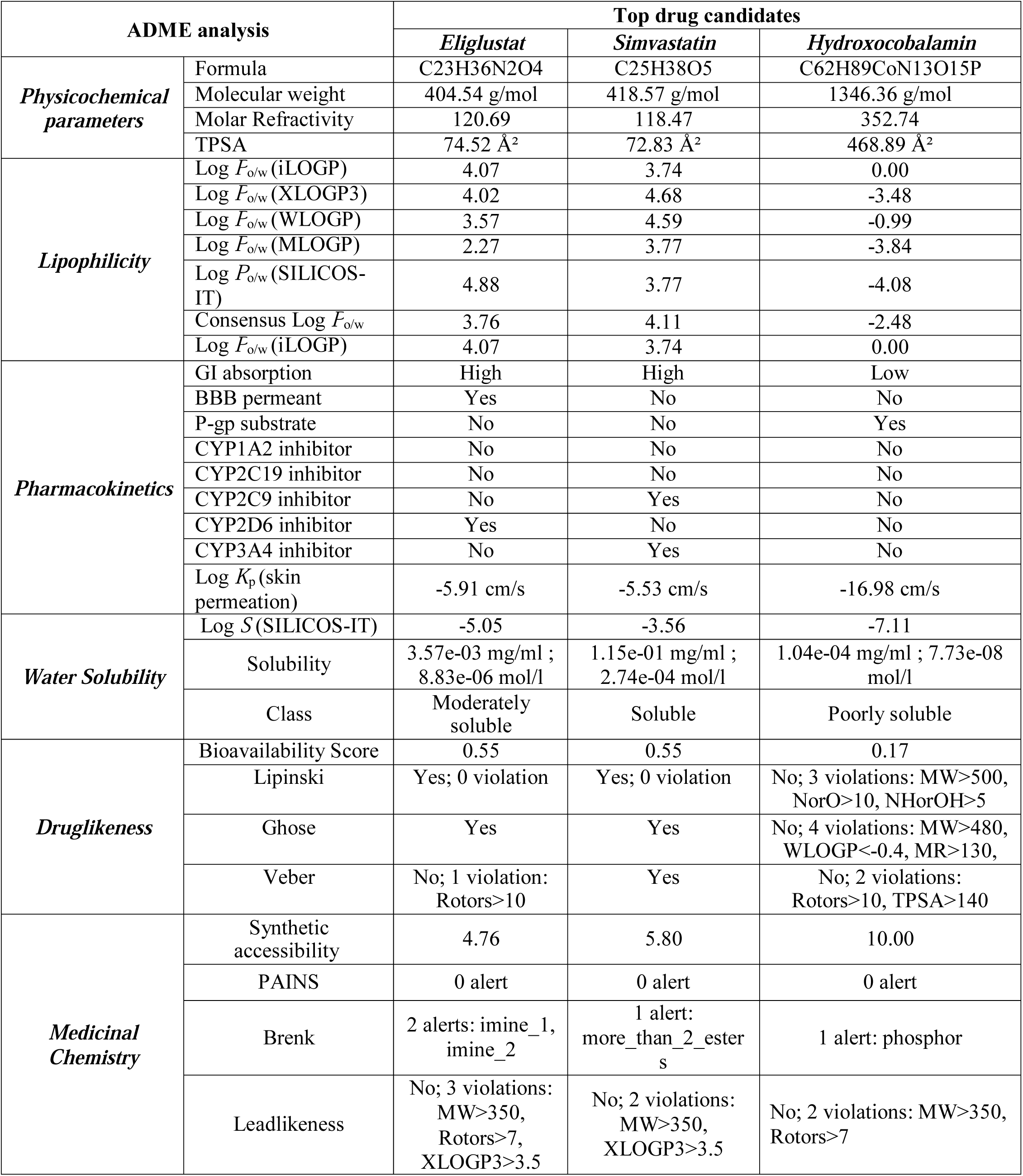
ADME profiling of top drug candidates.

### Screening of novel outer membrane proteins for vaccine construction

From the 25 novel outer membrane proteins (Supplementary file 14) designated as Set 11, two were selected based on the highest antigenicity score and human microbiome analysis to develop a novel chimeric peptide vaccine against *V. parahemolyticus* (Table 6). The schematic diagram summarizing the protocol over *in silico* vaccinomics strategy has been elucidated in Fig. 1. Sensor histidine protein kinase (Q87HJ8) and flagellar hook-associated protein (Q87JH9) possessed better antigenicity (0.64 and 0.53 respectively) while showed less percentage similarity (<45% and <41% respectively) when compared with gut microbiome data (Supplementary Table 4).

**Table 6:** Novel vaccine targets proteins showing higher antigenicity

| Accession No. | Protein name | VaxiJen score | Similarity with human microbiome (%) |
| --- | --- | --- | --- |
| Q87HJ8 | Sensor histidine protein kinase UhpB | 0.65 | <45 |
| Q87JH9 | Flagellar hook-associated protein | 0.53 | <41 |

### T-cell epitope prediction, transmembrane topology screening and antigenicity analysis

A plethora of CTL and HTL epitopes were identified for both proteins that can bind to the different large number of HLA-A and HLA-B alleles using MHC class-I and MHC class II binding predictions of IEDB (Supplementary File 15). Top epitopes (MHC-I and MHC-II binding peptides) for both proteins having the capacity to elicit strong T-cell responses were selected as putative T cell epitope candidates according to their topology screening by TMHMM and antigenic scoring (AS) by Vaxijen server (Supplementary File 16).

### Population coverage, allergenicity, toxicity and conservancy analysis

Two different population coverages were calculated from CTL and HTL populations for MHC class I and MHC class II restricted peptides, respectively (Fig. 4). Epitopes, found to be non-allergen for humans, were identified according to the allergenicity assessment via four servers (Supplementary File 17). However, epitopes predicted as a toxin was removed from the proposed list of epitopes. Several epitope candidates from both proteins were found to be highly conserved within different strains of *V. parahemolyticus* with maximum conservancy level of 99% for histidine protein kinase and 100% for flagellar hook-associated protein respectively (Supplementary Table 5). Top 3 epitopes (CTL and HTL) for each protein were considered based on the above-mentioned parameters to design the final vaccine construct (Table 7).

**Table 7:** Predicted final CTL and HTL epitopes of histidine protein kinase and flagellar hook-associated protein

| Protein | MHC | Epitope | Start | End | Vaxijen | Conservancy |
| --- | --- | --- | --- | --- | --- | --- |
| Sensor histidine protein kinase UhpB | MHC-I | AILLFPFAL | 36 | 44 | 2.993 | 99.00% |
|  |  | ILLFPFALR | 37 | 45 | 2.9721 | 99.00% |
|  |  | HDDGVGFKV | 448 | 456 | 2.363 | 99.00% |
|  | MHC-II | ILLFPFALRLGIALH | 37 | 51 | 2.1484 | 87.00% |
|  |  | AILLFPFALRLGIAL | 36 | 50 | 1.9451 | 87.00% |
|  |  | LLFPFALRLGIALHT | 38 | 52 | 1.8268 | 87.00% |
| Flagellar hook-associated protein | MHC-I | FNAQDEEGH | 125 | 133 | 1.8425 | 98.00% |
|  |  | GGRHNNLDL | 234 | 242 | 1.7855 | 99.00% |
|  |  | KPSPNFQAEV | 203 | 212 | 1.4978 | 45.00% |
|  | MHC-II | DSIESSFNAQDEEGH | 199 | 133 | 1.304 | 88.00% |
|  |  | IGGRHNNLDLMDGAH | 233 | 247 | 0.801 | 59.00% |
|  |  | KLSDDPMASIKLLNL | 38 | 52 | 0.887 | 88.00% |

**Figure 4.**
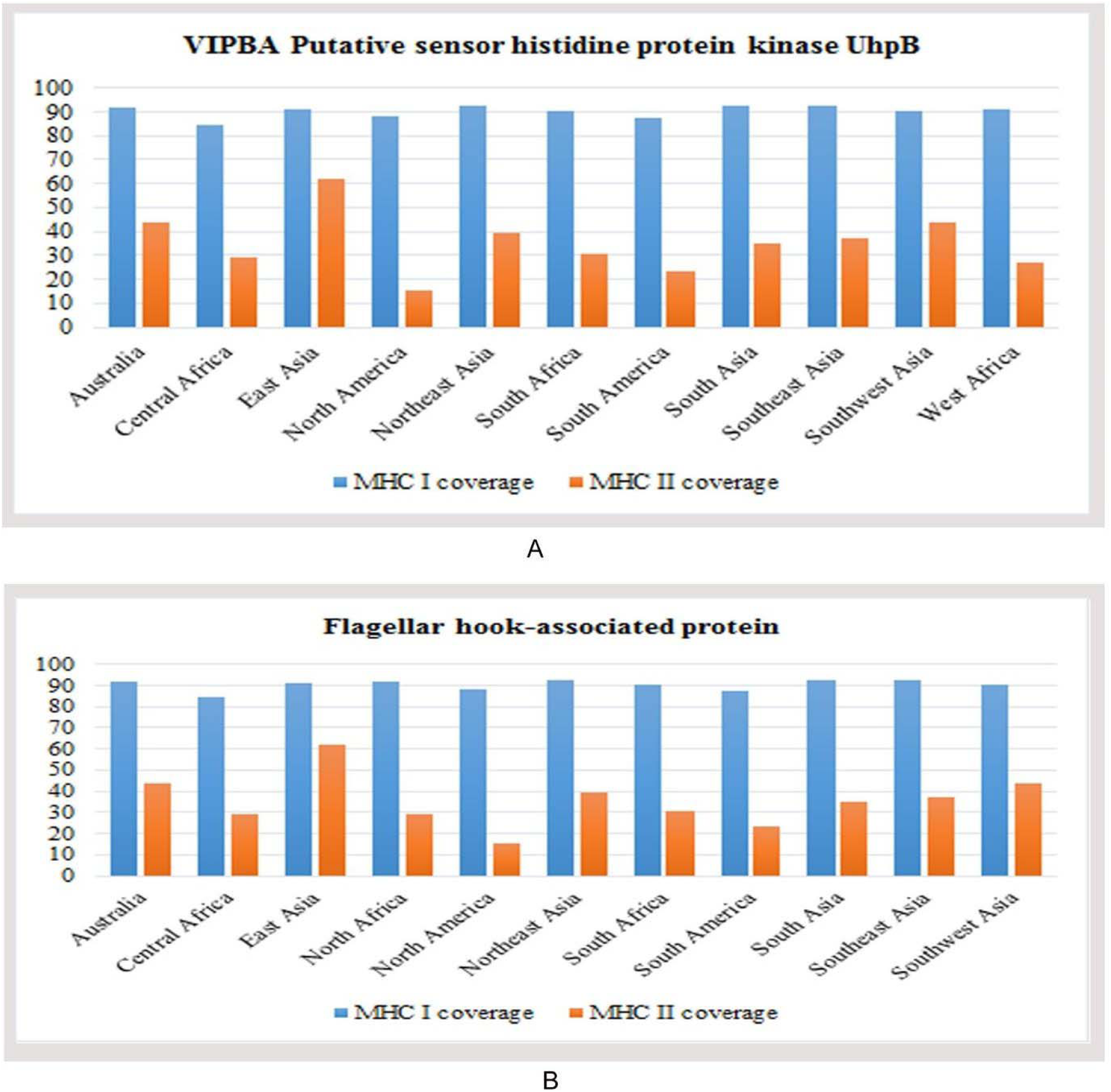
Population coverage analysis of (A) VIPBA putative sensor histidine protein kinase UhpB, and (B) Flagellar hook-associated protein.

### Prediction of 3D structures for superior epitopes and analysis of molecular docking

The epitopes, showing conservancy pattern at a biologically significant level, were only allowed for further docking analysis. 3D structures were predicted for top epitopes (6 from Flagellar hook-associated protein and six from sensor histidine protein kinase) to analyze their interactions with different HLA alleles. The PEP-FOLD3 server modeled five 3D structures for each epitope, and the best one was identified for docking study. The result showed that ‘AILLFPFAL’ epitope of Flagellar hook-associated protein was superior in terms of free binding energy while interacted with HLA-A* 11:01 (−8.3 kcal/mol). Demonstrated energy was −9.1 kcal/mol for epitope ‘GGRHNNLDL’ of Sensor histidine protein kinase (Table 8).

**Table 8:** Binding energy of predicted epitopes with selected MHC class I and MHC class II molecules generated from molecular docking by AutoDock

| Protein | Epitope | MHC Class | Binding Energy |
| --- | --- | --- | --- |
| Sensor histidine protein kinase | FNAQDEEGH | HLA-A*11:01 | -8.6 |
|  | GGRHNNLDL |  | -9.1 |
|  | KPSPNFQAEV |  | -8.2 |
|  | DSIESSFNAQDEEGH | HLA-DRB1*04:01 | -7.3 |
|  | IGGRHNNLDLMDGAH |  | -6.5 |
|  | KLSDDPMASIKLLNL |  | -6.3 |
| Flagellar hook-associated protein | AILLFPFAL | HLA-A*11:01 | -8.3 |
|  | ILLFPFALR |  | -8.1 |
|  | HDDGVGFKV |  | -8.1 |
|  | ILLFPFALRLGIALH | HLA-DRB1*04:01 | -7.0 |
|  | AILLFPFALRLGIAL |  | -6.4 |
|  | LLFPFALRLGIALHT |  | -7.2 |

### Identification of B-Cell epitope

B-cell epitopes of both proteins were generated using six different algorithms from IEDB (Supplementary Fig. 1 and Supplementary Fig. 2). The epitopes were further investigated to reveal their non-allergenicity pattern, and the top one epitope from each prediction was selected for vaccine construction (Supplementary Table 6).

### Epitope cluster analysis and vaccine construction

The construction of vaccine protein was based on identifying larger cassettes containing multiple epitopes. Epitope cluster analysis tool from IEDB predicted 21 epitope clusters among the top epitopes (6 CTL, 6 HTL epitopes, and 12 BCL epitopes) proposed in table 8 and supplementary table 2. Each vaccine construct was occupied by a protein adjuvant, PADRE peptide sequence, T-cell and B-cell epitopes with their respective linkers (Supplementary Table 7). Constructs V1, V2 and V3 were 370, 455 and 484 residues long, respectively. PADRE sequence was used to enhance the potency and efficacy of the peptide vaccine.

### Allergenicity, antigenicity and solubility prediction of different vaccine constructs

Results revealed V1 as the most potent vaccine candidate with better antigenic nature (1.18) and non-allergic behavior that can elicit a strong immune response (Supplementary Table 7). All three constructs showed solubility above the threshold value (0.45). Again, construct V1 was superior in terms of solubility potential. The surface distribution of charge, hydrophobicity and stability were calculated at 91 different combinations of pH and ionic strength (Fig. 5).

**Figure 5.**
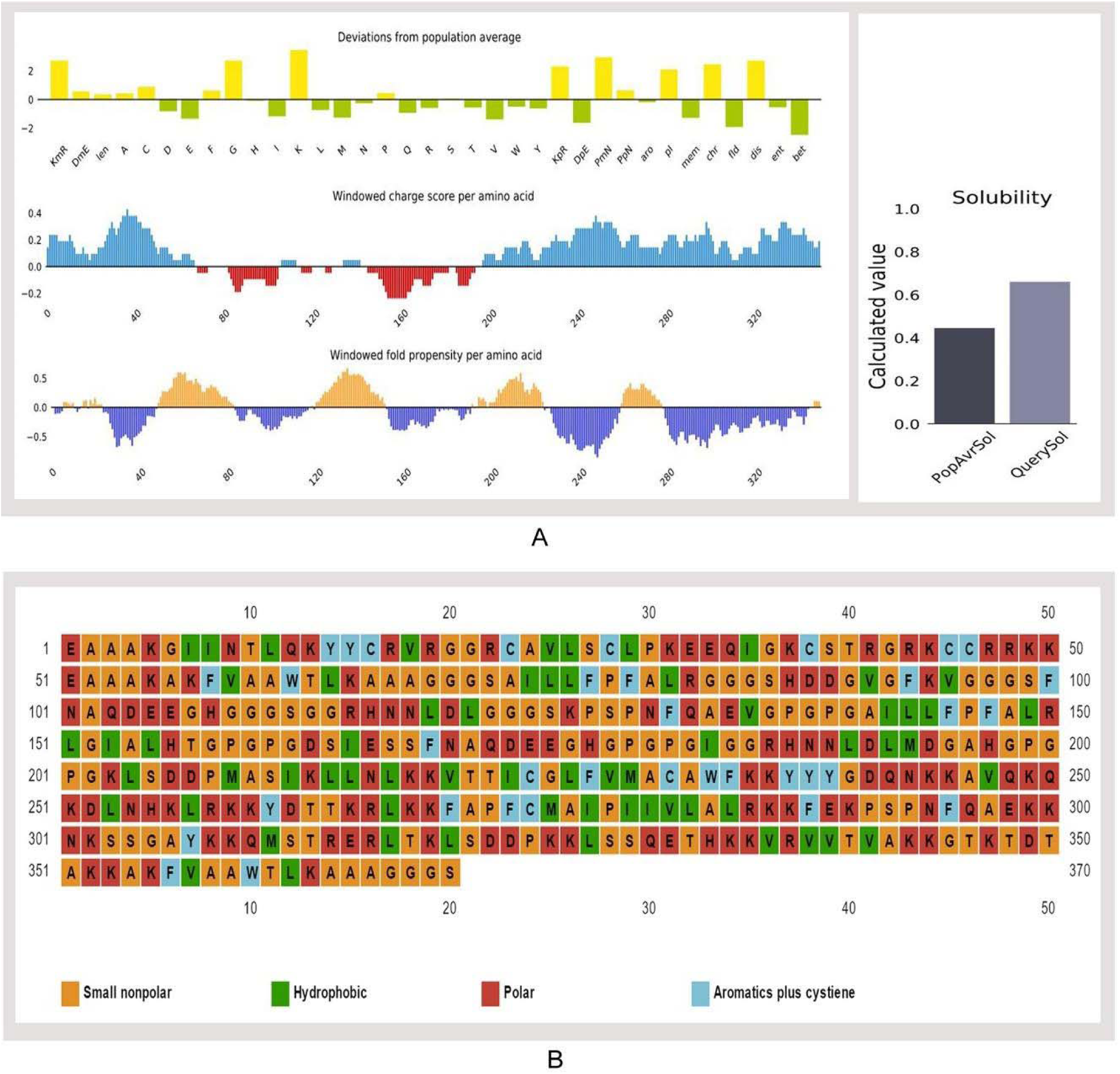
Solubility prediction of vaccine constructs. (A) Solubility prediction of designed vaccine construct V1 using via Protein-sol server, and (B) prediction of polar, nonpolar, hydrophobic and aromatic regions.

### Physicochemical characterization and secondary structure analysis

The molecular weight of the designed construct V1 was 39.45 kDa. The theoretical pI 9.95 implied that the protein would have a net negative charge above this pI and vice versa. At 0.1% absorption, the extinction coefficient was 26,930, while assuming all cysteine residues reduced. The estimated half-life was predicted to be >10 h in *E. coli* in vivo, whereas 1 h within mammalian reticulocytes in vitro. Hydrophilic behavior and thermostability of the protein were represented by the GRAVY value and aliphatic index that were−0.510 and 67.62, respectively. Instability index (37.49) and various physicochemical features classified the protein as a stable one with the capacity to induce a robust immunogenic reaction in the body. The predicted secondary structure confirmed to have 35.6% alpha helix, 11.89% sheet and 52.43% coil region (SupplementaryFig. 3). Around 34.59% polar, 16.21% hydrophobic and 9.46% aromatic regions were identified in the structure (Fig. 5).

### Tertiary structure prediction, refinement, validation and disulfide engineering of vaccine construct

I-TASSER predicted five models for each proposed vaccine candidates, which were ranked based on cluster size. Ten best templates (with the highest Z-score) selected from the LOMETS threading programs were used to predict the tertiary structures. Homology modeling was performed by using 1kj6 from RCSB Protein Data Bank) as a best suited template for Vaccine protein V1. Results showed that model 1 had the highest C-Score of −2.11 while the estimated TM-score and RMSD were 0.46±0.15 and 11.6±4.5Å (Fig. 6). After refinement, 88.3% and 98.1% residues were in the favored and allowed region revealed by Ramachandran plot analysis (Fig. 6). The modeled tertiary structure of designed construct V2 and V3 have been shown in Supplementary Fig. 4. A total of 22 pairs of amino acid residues were identified with the potential to form disulfide bonds by DbD2 server. However, only two pairs (i.e., ARG 82-Gly 85, Lys 347-Thr 350) were compatible with disulfide bond formation considering the energy, chi3 and B-factor parameter (Supplementary Fig. 5). All these residues were replaced with a cysteine residue.

**Figure 6.**
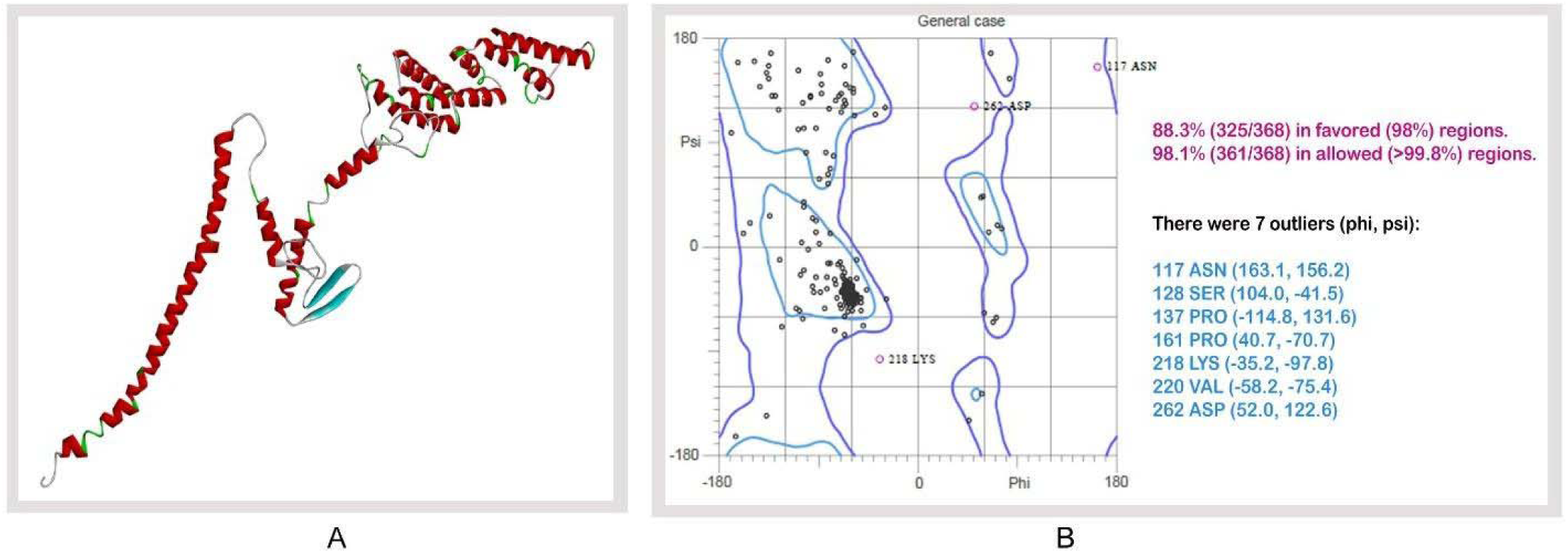
Tertiary structure prediction and validation of vaccine protein V1. (A) Tertiary structure of modeled construct V1, (B) Ramachandran plot analysis of vaccine protein V1.

### Protein-protein docking and molecular dynamics simulation

Docking study was conducted between three vaccine constructs (i.e., V1, V2, V3) and different HLA alleles. Construct V1 showed biologically significant results and found to be superior in terms of free binding energy (Supplementary Table 8). Besides, the binding affinity of the predicted vaccine and TLR1-2 heterodimer complex was also analyzed. The 3D structure of human TLR-1/2 heterodimer was retrieved from the RCSB protein data bank. ClusPro generated thirty protein-ligand complexes (clusters) as output along with respective free binding energy. The lowest energy was −1257.9 for cluster 1 (Fig. 7). FireDock output refinement of the PatchDock server showed the lowest global energy of −7.08 for solution 5. Normal mode analysis allowed the demonstration of large scale mobility and the stability of proteins. The analysis was performed based on the internal coordinates of the protein-protein complex. In the 3D model, the direction of each residue was given by arrows, and the length of the line represented the extent of mobility (Fig. 8A). The eigenvalue found for the complex was 2.4784e−05 (Fig. 8B). The vaccine protein V1 and TLR1-2 heterodimers were oriented towards each other. The B-factor values deduced from normal mode analysis was analogous to RMS (Fig. 8C). Hinges in the chain indicated the probable deformability of the complex measured by the contortion of each residue (Fig. 8D). The variance associated with each normal mode was inversely linked to the eigenvalue. Covariance matrix explained the coupling between pairs of residues was correlated, uncorrelated, or anti-correlated motions were represented via red, white and blue colors, respectively (Fig. 8E). The result also generated an elastic network model (Fig. 8F) that identified the pairs of atoms connected via springs. Each dot in the diagram was indicated one spring between the corresponding pair of atoms and colored based on the degree of stiffness.

**Figure 7.**
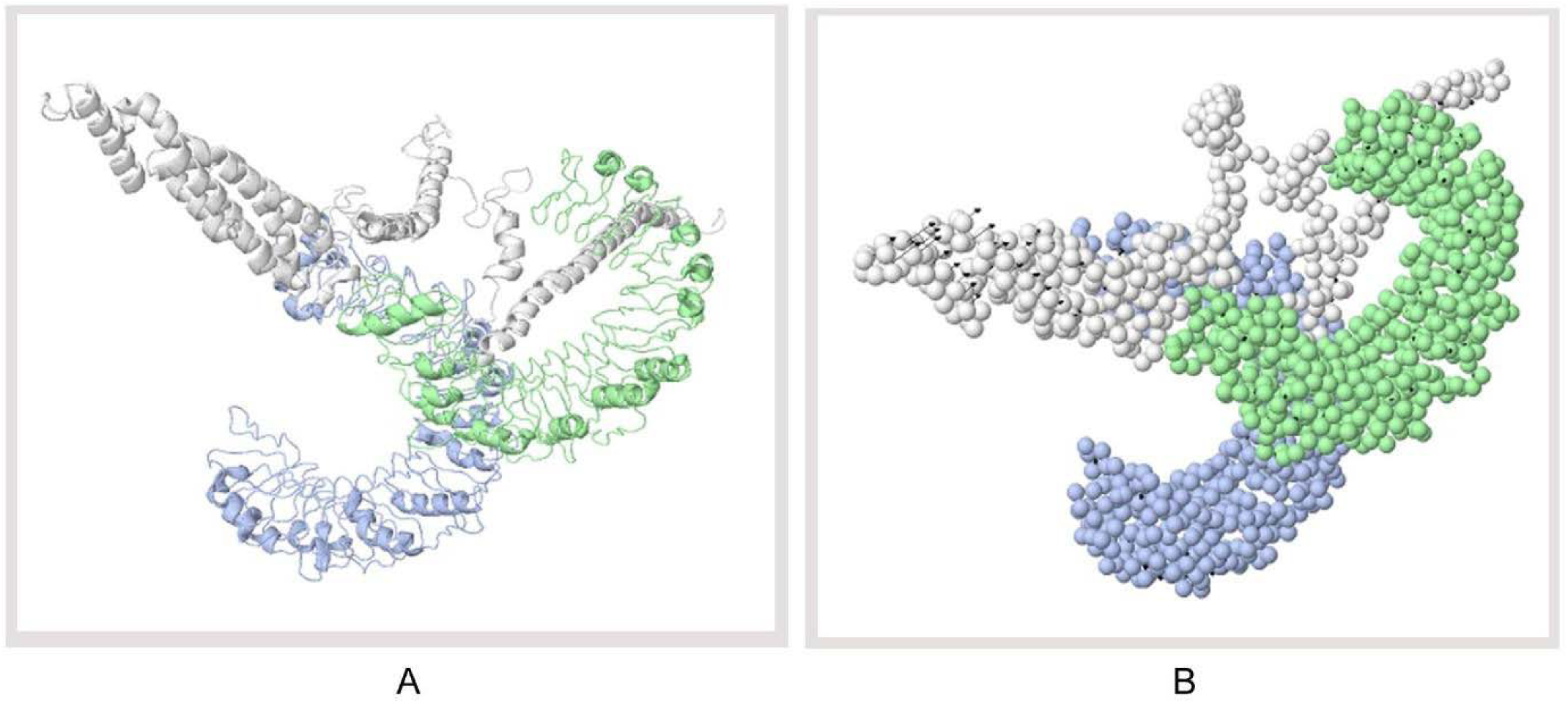
Docked complex of vaccine construct V1 with human TLR 1/2 heterodimer. (A) Cartoon format, and (B) Ball structure.

**Figure 8.**
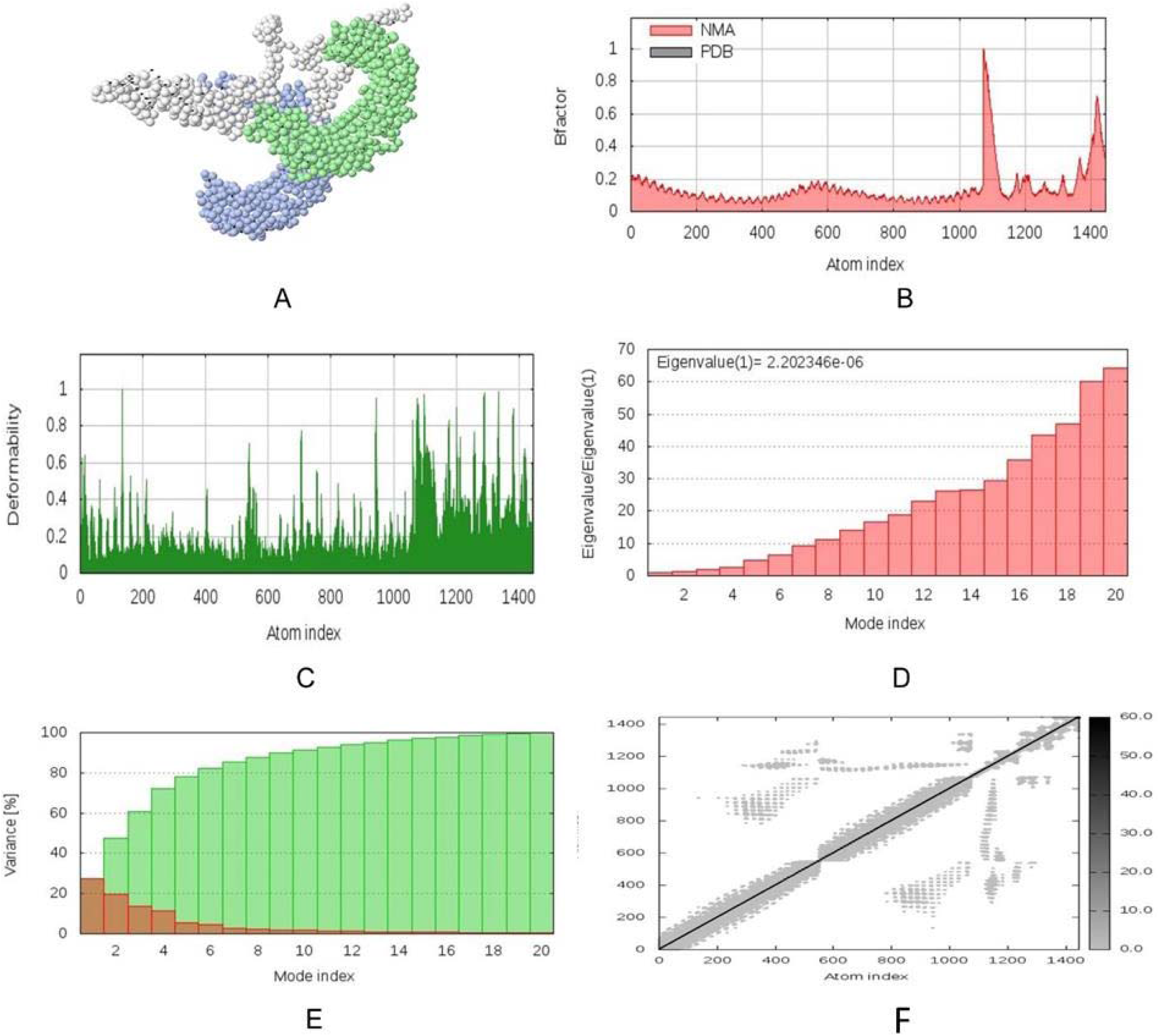
Molecular dynamics simulation of vaccine protein V1-TLR8 complex. Stability of the protein-protein complex was investigated through (A) mobility, (B) B-factor, (C) deformability, (D) eigenvalue, (E) covariance and (F) elastic network analysis.

### Codon adaptation, in silico cloning and similarity analysis with human proteins

*E. coli* strain K12 was selected as the host for the cloning purpose of the vaccine construct V1. Vaccine protein V1 was transcribed reversely, where the Codon Adaptation Index (CAI) was found 0.994, and the GC content of the optimized codons (50.55%) was also significant. The construct did not hold restriction sites for ApaI and BglI, which ensured its safety for cloning purposes. The optimized codons were incorporated into the pET28a(+) vector along with ApaI and BglI restriction sites. A clone of 5634 base pair was obtained, including the 1118 bp desired sequence and the rest belonging to the vector. The desired region was shown in red color in between the pET28a(+) vector sequence (Fig. 9). Sequence similarity analysis of the proposed vaccine with human proteins revealed that there was no similarity between predicted vaccine constructs and human proteins.

**Figure 9.**
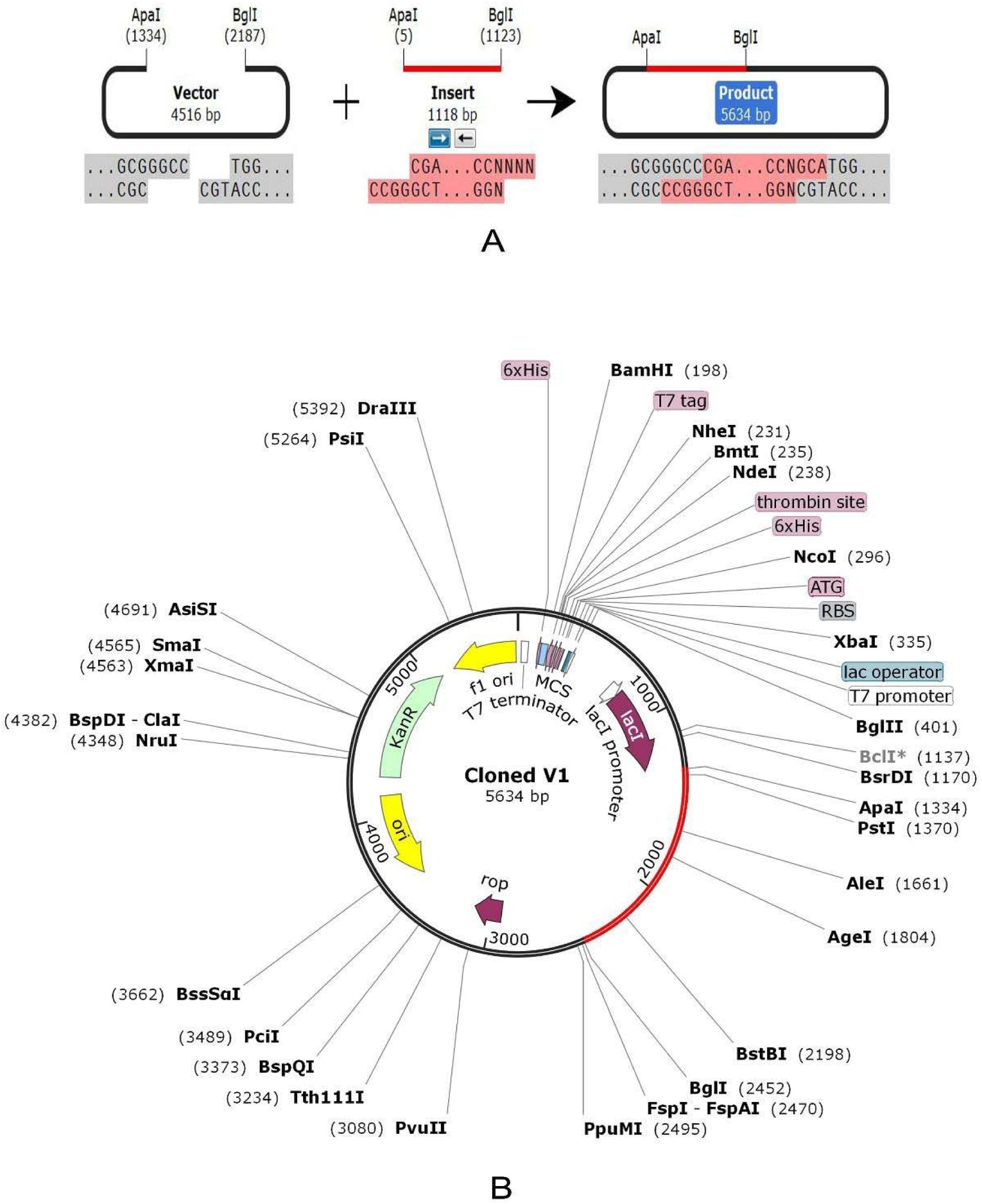
In silico restriction cloning of the gene sequence of construct V1 into pET28 a(+) expression vector; (A) Codon adaptation (B) Restriction digestion of the vector pET28 a(+) and construct V1 with BglI and ApaI (C) Inserted desired fragment (V1 Construct) between ApaI (1334) and BglI (2452) indicated in red color.

## Discussion

Due to severe effects on human health (Velazquez-Roman et al., 2014; Wu et al., 2014), the emergence of rapid antibiotic resistance (Golkar et al., 2014) and also having economic importance for substantially impairing the aquaculture production (Tang et al., 2014), it has become an urgent necessity to develop effective drug targets and vaccine candidates against *Vibrio parahaemolyticus.* Different computational approaches are now being widely practiced to identify proteins those are essential for the survival of the pathogen and not involved in the metabolic pathways of the host, thereby choosing the proteins associated only in the metabolic pathways of the pathogen is important (Judson and Mekalanos, 2000; Hossain et al., 2013). Essential proteins are most promising for new antibacterial drug targets since most antibiotics are designed to bind essential gene products (Zhang et al., 2004). Here, subtractive genome approaches (removal of paralogous proteins, identification of non-homologous proteins against the host, identification of essential proteins and metabolic pathways analysis of the pathogen), and vaccinomics strategy were employed for identifying novel drug and vaccine molecules through the comprehensive proteome exploration of *Vibrio parahaemolyticus* genome.

The complete proteome of *V. parahaemolyticus* (4822 proteins) was retrieved from the NCBI database, and the homologous proteins were removed based on their identity with human proteins. Proteins encoded by essential genes and unique to an organism can be considered as species-specific drug targets (Mondal et al., 2015), as they play vital roles in its metabolism. The study revealed 96 unique, essential proteins (Set5) of *V. parahaemolyticus*, which can be considered as suitable drug targets for combating *V. parahaemolyticus* infections. Among the unique proteins, 55 proteins were druggable and can be targeted using existing drugs (92) that are already approved and available in the market (Supplementary Table 1). In the case of a broad-spectrum drug, for avoiding mutational changes as well as the emergence of resistant bacteria, the DrugBank databases were screened, which contains entries for 2556 approved small molecule drugs, 962 approved biotech drugs, 112 nutraceuticals and over 5125 experimental drugs. A total of 41 proteins of *V. parahaemolyticus* showed no similarity after passing through the DrugBank database and listed for the prediction of novel drug targets and vaccine candidates (set 6). To avoid severe cross-reaction and toxic effects in human, identification of nonhomologous proteins to essential human proteins (referred to as ‘anti-targets’) was a crucial step considered in this study. However, the identified novel drug targets (41) showed no evidence of similarity with the ‘anti-targets’. Although both cytoplasmic and membrane proteins serve the purpose as therapeutic targets (Michael et al., 2014), membrane proteins are best suited for vaccine candidates (Baliga et al., 2018; Hasan et al., 2019a). Hence, in this study, membrane proteins (25) were used for vaccine construction, whereas cytoplasmic proteins (16) were proposed as suitable drug targets. Targeting human microbiome non-homology proteins is suitable since drugs or vaccines designed and administered for these targets will be less harmful to other commensal microbial strains. Among the novel cytoplasmic proteins (16), only the proteins (9) conferring <45% similarity with human microbiota were retained. TheVFDB (virulence factor database) analysis studies confirmed that ‘VIBPA Type II secretion system protein L (Q87TC9)’ and ‘VIBPA Putative fimbrial protein Z (Q87I65)’ were associated with virulence in the host (Set 10). The protein-protein interaction studies also strengthened the superiority of these two proteins as suitable drug targets (Table 4).

Molecular docking of 350 human metabolites against ‘VIBPA Type II secretion system protein L’ and ‘VIBPA Putative fimbrial protein Z’ was conducted to screen superior drug molecules (Supplementary Table 2). The study revealed that ‘Eliglustat (DB09039)’ was the top drug molecules for both protein targets in terms of free binding energy (Table 5). Therefore, it can be suggested as a suitable drug to treat infections caused by *V. parahaemolyticus*. Moreover, drug profiling (Physicochemical parameters, Lipophilicity, Pharmacokinetics, Water Solubility, Druglikeness, Medicinal Chemistry) of ‘Eliglustat (DB09039)’ along with other top candidates, i.e., ‘Simvastatin (DB09039)’ and ‘Hydroxocobalamin (DB00200)’ were also performed through ADME analysis (Table 5).

Several advantages help the researchers to select membrane proteins both as drug and vaccine candidates as their functions can be easily studied through computer-based approaches than wet-lab process (Hasan et al., 2019a; Hasan et al., 2019b). In this study, two vaccine targets, ‘Sensor histidine protein kinase UhpB (Q87HJ8)’ and ‘Flagellar hook-associated protein (Q87JH9)’ were selected after screening the novel outer-membrane proteins (25) based on their antigenicity score and human microbiome non-homology analysis (Table 6). Both the proteins further analyzed to design a potent, highly immunogenic vaccine candidate against *V. parahaemolyticus.* Numerous antigenic epitopes were generated which were investigated extensively for antigenicity, allergenicity, toxicity, conservancy and other physiochemical properties using a number of bioinformatics tools and software. The final vaccine constructs were designed with the help of different adjuvants and amino acid linkers (Solanki and Tiwari, 2018). It has been reported that the PADRE sequence reduces the polymorphism of HLA molecules in the population (Ghaffari-Nazari et al., 2013; Hasan et al., 2019b). Linkers in vaccines also enhanced the immunogenicity of the vaccines in previous studies (Yang et al., 2015; Hasan et al., 2019c). Therefore, all the important that could induce the immunogenicity of the designed vaccine constructs were taken. Also, disulfide engineering was employed to enhance the stability of the designed vaccine. The purpose of the molecular docking analysis was to show the proposed epitopes could interact with at least one MHC molecule at minimum binding energy. Therefore, it was done to explore the binding affinity of promiscuous epitopes with different HLA alleles including HLA-DRB1*03:01 (1A6A), (HLA-DRB5*01:01 (1H15), HLA-DRB3*01:01 (2Q6W), HLA-DRB1*04:01 (2SEB), HLA-DRB1*01:01 (2FSE), and HLA-DRB3*02:02 (3C5J). The OmpU, one of the major outer membrane porins of *V. parahaemolyticus*, is recognized by the Toll-like receptor 1/2 (TLR1/2) heterodimer in THP-1 monocytes (Gulati et al., 2019). So, a docking study was also performed to analyze the affinity between the designed construct and human TLR1/2 heterodimer. The vaccine receptor complex showed deformability at a minimum level, which also strengthened our prediction. Finally, the optimized codons of the designed construct been cloned into the pET28a(+) vector of *E. coli* strain K12.

The idea of subtractive genomic analysis using various bioinformatics tools has brought a revolution in the drug discovery process. The present study will help to develop novel therapeutics and preventive measures against *V. parahaemolyticus*, thereby help to reduce the mortality and morbidity caused by it. However, further in vivo trials using model organisms are highly recommended to validate our prediction.

## Materials and Methods

The whole proteome of *V. parahemolyticus* was analyzed according to subtractive genomic approach to recognize novel drug targets as well as vaccine candidates. The overall workflow for subtractive proteomic analysis and vaccinomics approach has been illustrated in Fig. 1 and Fig. 2, respectively.

### Retrieval of complete proteome and identification of essential proteins

The whole proteome of *V. parahemolyticus* strain O3:K6 was retrieved from NCBI Genome database. Paralogous sequences were excluded from the proteome of *V. parahemolyticus* by using CD-HIT (Li and Godzik, 2006). With a cutoff score of 0.6, proteins with more than 60% identity were excluded. Remaining proteins were subjected to BLASTp against *H. sapiens* human Refseq proteome in ‘Ensemble Genome Database 92’ using threshold expectation value (E-value) 10^−3^ as the parameter. Proteins were assumed as homologous were excluded if any significant hit above the threshold value 10^−4^ was found. The remaining non-homologous proteins were subjected to the Database of Essential Genes (DEG) (Zhang et al., 2004; Luo, 2013). Proteins hit with expectation value ≤10-100, identity ≥ 25% were listed for metabolic pathway analysis considering as essential non-homologous proteins of *V. parahemolyticus*.

### Analysis of metabolic pathways

Kyoto Encyclopedia of Genes and Genomes which contains complete metabolic pathways present in living organisms (Moriya et al., 2007). Metabolic pathways of *V. parahemolyticus* were analyzed against the human metabolic pathways through the KEGG server. All metabolic pathways present in the pathogen (*V. parahemolyticus*) and host (*H. sapiens*) were collected from the KEGG PATHWAY database using three letters KEGG organism code ‘vpa’ and ‘has’ respectively. A comparison was made in order to recognize the unique metabolic pathways only present in the *V. parahemolyticus*, while the remaining pathways of the pathogen were grouped as a common one. Identified host non-homologous, essential proteins of *V. parahemolyticus* were subjected to BLASTp through the KAAS server at KEEG. Proteins present only in the unique metabolic pathways of the pathogen were listed for further analysis.

### Druggability analysis and identification of novel drug targets

A ‘druggable’ target needs to have the potentiality to bind to the drugs and drug-like molecules with high affinity. Shortlisted unique proteins were screened through the database of DrugBank 5.1.0 (Wishart et al., 2017) using default parameters to identify both druggable proteins and novel therapeutic targets.

### ‘Anti-target’ analysis and prediction of subcellular localization

This analysis was performed to avoid any kind of cross-reactivity, and toxic effects due to docking between the drugs administered for the pathogen and the host ‘anti-targets’.’Anti-targets’ are gene products that show cross-reactivity with administered therapeutics. Novel drug targets were subjected to BLASTp analysis in the NCBI blast program against these human ‘anti-targets’ setting an E-value <0.005, query length >30%, identity <25% as parameters. Proteins were showing a <25% identity that was listed for subcellular localization analysis. Besides, proteins functioning in cytoplasm can be used as putative drug targets, while surface membrane proteins can be considered both as drug targets and vaccine candidates. PSORTb v3.0.2 (http://www.psort.org/psortb/index.html), CELLO v.2.5 (http://cello.life.nctu.edu.tw/), ngLOC servers were used to predict subcellular localization of shortlisted pathogen-specific essential proteins.

### Human microbiome non-homology analysis

Both membrane and cytoplasmic proteins were subjected to BLASTp through NCBI protein blast server against the dataset present in the Human Microbiome Project server (http://www.hmpdacc.org/hmp/) “(BioProject-43021)” (Turnbaugh et al., 2007) with a cutoff score 0.005. Membrane proteins showing <45% similarity were selected for vaccine candidacy, whereas cytoplasmic proteins showing <45% similarity were selected for protein-protein interaction analysis.

### Analysis of virulence factors (VF’s) and protein-protein interactions studies (PPIs)

Virulence factors are responsible for modulating or degrading host defense mechanisms by bacteria. Novel cytoplasmic proteins with the least similarity with the human microbiome were subjected to BLASTp search against the database of protein sequences from the VFDB core dataset (Chen et al., 2005) with default hit with cut-off bit score >100, and E-value was 0.0001. The protein-protein interactions studies (PPIs) of selected shortlisted proteins were predicted using STRING v10.5 (Szklarczyk et al., 2018). PPIs with a high confidence score (≥90%) were considered to avoid false-positive results. Only characterized proteins were subjected to BLASTp.

### Screening of drug molecules against novel cytoplasmic proteins

All the pharmaco-metabolites reported in the Human Metabolites Database (www.hmdb.ca) were used for the screening of suitable drugs. Molecular docking was performed against predicted drug targets (novel cytoplasmic proteins) by using AutoDock Vina tools (Trott and Olson, 2010). The size of the grid box was set to 54 A° x 74 A° x 126 A° (x, y and z) and 65A° x 85 A° x 65 A° (x, y and z) with 1 A° spacing between the grid points for two cytoplasmic therapeutic target proteins (Q87TC9 and Q87165 respectively).

### Screening of novel outer membrane proteins for vaccine construction

The VaxiJen v2.0 (http://www.ddg-pharmfac.net/vaxijen/) was used for the investigation of protein immunogenicity to find the most potent antigenic outer membrane proteins (Doytchinova and Flower, 2007a). Proteins were prioritized based on their antigenic score (threshold value 0.4) and sequence similarity with human microbiota.

### T-cell epitope prediction, transmembrane topology screening and antigenicity analysis

MHC-I (http://tools.iedb.org/mhci/) and MHC-II prediction tool (http://tools.iedb.org/mhcii/) prediction tool of the Immune Epitope Database were used to predict the MHC-I binding and MHC-II binding peptides, respectively. To predict the transmembrane helices in proteins (Krogh et al., 2001) and to determine epitope antigenicity (Doytchinova and Flower, 2007b), TMHMM (http://www.cbs.dtu.dk/services/TMHMM/) and VaxiJen v2.0 server (http://www.ddg-pharmfac.net/vaxijen/) were utilized.

### Population coverage, allergenicity, toxicity and conservancy analysis

Population coverage for each epitope was analyzed by the IEDB population coverage calculation tool (http://tools.iedb.org/population/) (Vita et al., 2014). The most potent antigenic epitopes were selected and allowed for determining the allergenicity pattern via four servers named AllergenFP (Dimitrov et al., 2014), AllerTOP (http://www.ddg-pharmfac.net/AllerTop/) (Dimitrov et al., 2013), Allermatch (http://www.allermatch.org/allermatch.py/form) (Fiers et al., 2004) and Allergen Online (Goodman et al., 2016). Moreover, the ToxinPred server predicted the toxicity level of the proposed epitopes (http://crdd.osdd.net/raghava/toxinpred/) (Gupta et al., 2013). The conservancy level determines the efficacy of epitope candidates to confer broad-spectrum immunity. For revealing the conservancy pattern, homologous sequence sets of the selected antigenic proteins were retrieved from the NCBI database by using the BLASTp tool. Further, the epitope conservancy analysis tool (http://tools.iedb.org/conservancy/) at IEDB was selected for the analysis of the conservancy pattern.

### Prediction of 3D structures for superior epitopes and analysis of molecular docking

Top-ranked epitopes were subjected to the PEP-FOLD server to predict peptide structures (Wang et al., 2011). Depending on the available structures deposited in Protein Data Bank (PDB) database, HLA-A*11:01 and HLA-DRB1*04:01 were selected for docking analysis with MHC class I and class II binding epitopes respectively. MGLTools were used to visualize and analyze the molecular structures of biological compounds (Sanner, 1999). The grid box was set to 28 A°, 18 A°, 20 A° (x, y and z) with a default value of 1.0 A° spacing by AutodockVina at 1.00-°A spacing. The exhaustiveness parameter was kept at 8.00, and the number of outputs was set at 10 (Morris et al., 2009). Output PDBQT files were converted in PDB format using Open Babel. The docking interaction was visualized with the PyMOL molecular graphics system (https://www.pymol.org/).

### Identification of B-Cell epitope

B cell epitopes were predicted for both proteins to find the potential antigens that would interact with B lymphocytes and initiate the immune response. Several tools from IEDB i.e. Kolaskar and Tongaonkar antigenicity scale (Kolaskar and Tongaonkar, 1990), Karplus and Schulz flexibility prediction (Karplus and Schulz, 1985), Bepipred linear epitope prediction analysis (Jespersen et al., 2017), Emini surface accessibility prediction (Emini et al., 1985), Parker hydrophilicity prediction (Parker et al., 1986) and Chou and Fasman beta-turn prediction (Chou and Fasman, 1978) were used to identify the B cell antigenicity depending on six different algorithms.

### Epitope cluster analysis and vaccine construction

Epitope cluster analysis tool from IEDB was used to identify the epitope clusters with overlapping peptides for both proteins using the top CTL, HTL and BCL epitopes as input. The identified clusters and singletons were further utilized to design construct. Vaccine sequences started with an adjuvant followed by the top CTL epitopes, top HTL epitopes and BCL epitopes respectively for both proteins. Three vaccine constructs i.e. V1, V2 and V3, each associated with different adjuvants including beta-defensin (a 45 mer peptide), L7/L12 ribosomal protein and HABA protein (*M. tuberculosis*, accession number: AGV15514.1) (Rana and Akhter, 2016) were constructed. PADRE sequence and different linkers, for instance, EAAAK, GGGS, GPGPG and KK linkers, were also used to construct effective vaccine molecules.

### Allergenicity, antigenicity and solubility prediction of different vaccine constructs

The AlgPred v.2.0 (Saha and Raghava, 2006) and AllerTOP v.2.0 (Dimitrov et al., 2013) servers were utilized to predict the non-allergic behavior of the vaccine constructs. For proposing the superior vaccine candidate, the VaxiJen v2.0 server (Doytchinova and Flower, 2007b) was utilized. The probable antigenicity of the constructs was determined through an alignment-independent algorithm. Protein-sol software (Hebditch et al., 2017) predicted the solubility score of the proposed vaccines.

### Physicochemical characterization and secondary structure analysis

The ProtParam, a tool from Expasy’s server (http://expasy.org/cgi-bin/protpraram)(Gasteiger et al., 2003; Hasan et al., 2015) was used to characterize the vaccine proteins functionally – including molecular weight, isoelectric pH, aliphatic index, hydropathicity, instability index, GRAVY values and estimated half-life and other physicochemical properties were investigated. The PSIPRED v3.3 (Kosciolek and Jones, 2014; Packer, 2012) were used to predict the alpha helix, beta sheet and coil structure of the vaccine protein. The polar, nonpolar and aromatic regions were also determined.

### Tertiary structure prediction, refinement, validation and disulfide engineering of vaccine construct

The I-TASER server (Yang et al., 2015) was employed for determining the 3D structure of designed vaccine constructs based on the degree of similarity between the target protein and available template structure from PDB. Refinement was performed using ModRefiner (Xu and Zhang, 2011). The refined protein structure was validated through the Ramachandran plot assessment by the MolProbity software (Chen, V.B. et al., 2010). Residues in the highly mobile region of the protein exhibit the potential to be mutated with cysteine. Pairs of residues with proper geometry and the ability to form a disulfide bond were detected by the DbD2 server to perform disulfide engineering (Craig and Dombkowski, 2013). The value of chi3 considered for the residue screening was between −87 to +97 while the energy considered was<2.5.

### Protein-protein docking and molecular dynamics simulation

The binding affinity of the vaccine constructs with different HLA alleles and human immune receptors, ClusPro 2.0 (Comeau et al., 2004), hdoc (Macalino et al., 2018) and PatchDock server (Schneidman-Duhovny et al., 2005) were applied. Desirable complexes were identified according to better electrostatic interaction and free binding energy following refinement via the FireDock server (Mashiach et al., 2008). The iMODS server was used to explain the collective motion of proteins via analysis of normal modes (NMA) in internal coordinates (Lopez-Blanco et al., 2014). Essential dynamics is a powerful tool and alternative to the costly atomistic simulation that can be compared to the normal modes of proteins to determine their stability (Aalten et al., 1997; Wuthrich et al., 1980; Cui and Bahar, 2007). The server predicted the direction and magnitude of the immanent motions of the complex in terms of deformability, eigenvalues, B-factors and covariance. The structural dynamics of the protein-protein complex was investigated (Prabhakar et al., 2016).

### Codon adaptation, in silico cloning and similarity analysis with human proteins

A codon adaptation tool (JCAT) was used to adapt the codon usage to the well-characterized prokaryotic organisms for accelerating the expression rate in it. Rho-independent transcription termination, prokaryote ribosome binding site and cleavage sites of restriction enzyme ApaI and BglI were avoided while using the server (Grote et al., 2005). The optimized sequence of vaccine protein V1 was reversed, followed by conjugation with ApaI and BglI restriction site at the N-terminal and C-terminal sites, respectively. SnapGene (Solanki and Tiwari, 2018) restriction cloning module was used to insert the adapted sequence into the pET28a(+) vector between ApaI (1334) and BglI (2452). At last, human sequence similarity analysis of the proposed vaccine with human proteins was done by using NCBI protein-protein Blast (https://blast.ncbi.nlm.nih.gov/Blast.cgi), and here blast was done against Homo sapiens (taxid: 9606) dataset.

## Funding

The authors received no direct funding for this research.

## Author contributions

Mahmudul Hasan, Conceptualization, Data curation, Formal analysis, Investigation, Visualization, Methodology, Manuscript-original draft, review and editing; Kazi Faizul Azim, Abdus Shukur Imran and Ishtiak Malique Chowdhury, Resources, Data curation, Formal analysis, Validation, Investigation, Visualization, Manuscript writing; Shah Rucksana Akhter Urme, Md Sorwer Alam Parvez and Md Tahsin Khan, Software, Formal analysis, Validation, Investigation, Visualization, Manuscript writing; Md Bashir Uddin and Syed Sayeem Uddin Ahmed, Conceptualization, Resources, Data curation, Software, Analysis, Validation, Visualization, Methodology, Supervision, Manuscript-original draft, review and editing.

## Declaration of interests

The authors declare that they have no known competing financial interests or personal relationships that could have appeared to influence the work reported in this paper.

## Data availability

All data generated and analysed during this study is included in the main manuscript or supplementary files.

## Supplementary Figures

**Supplementary Figure 1.**
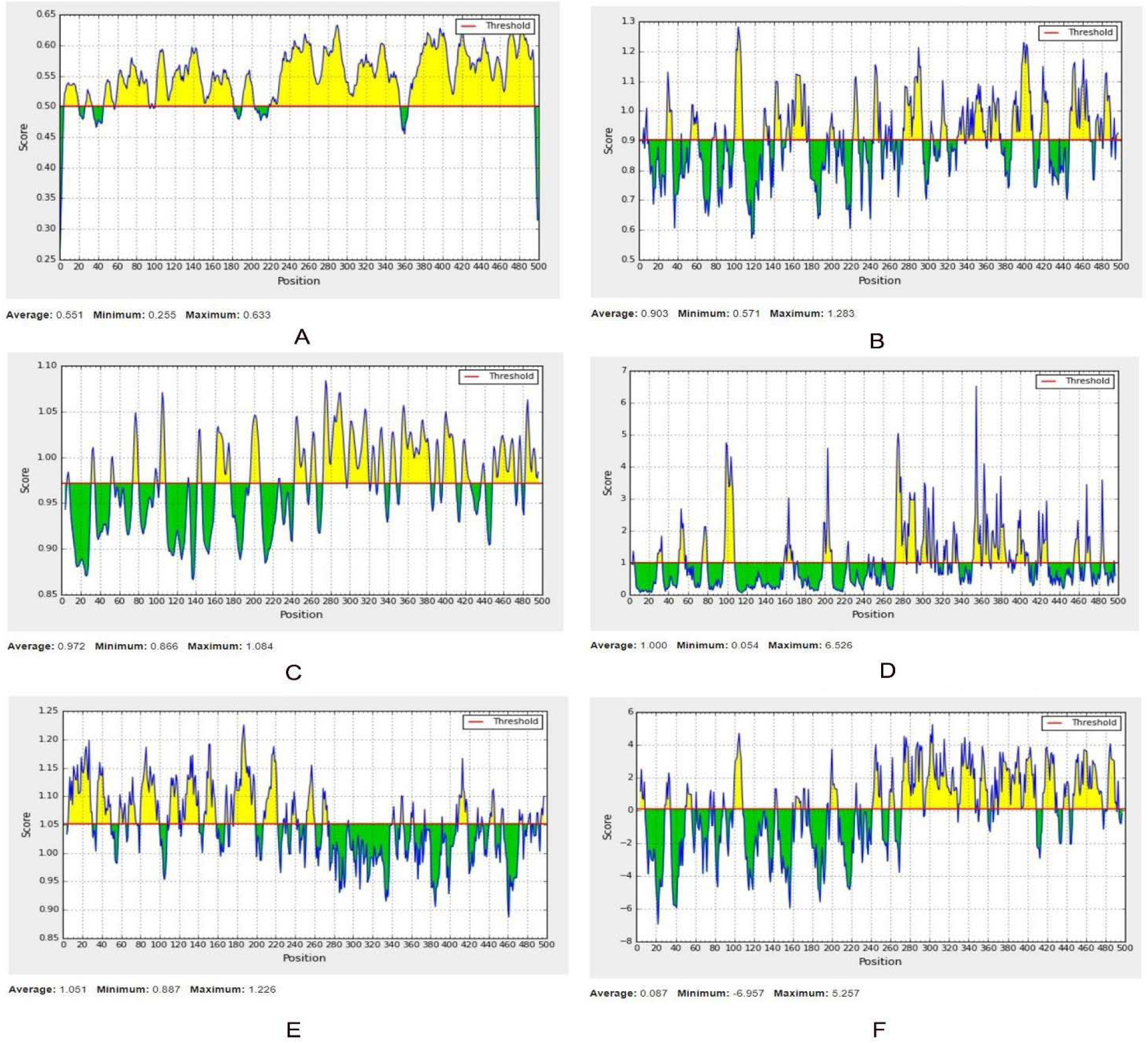
B-cell epitope prediction of Sensor histidine protein kinase UhpB (A: Linear, B: Beta-turn, C: Flexibility, D: Surface Accessibility, E: Antigenicity, F: Hydrophilicity). For each graph, X-axis and Y-axis represent the position and score. Residues that fall above the threshold value are shown in yellow color while the highest peak in yellow color identifies most favored position.

**Supplementary Figure 2.**
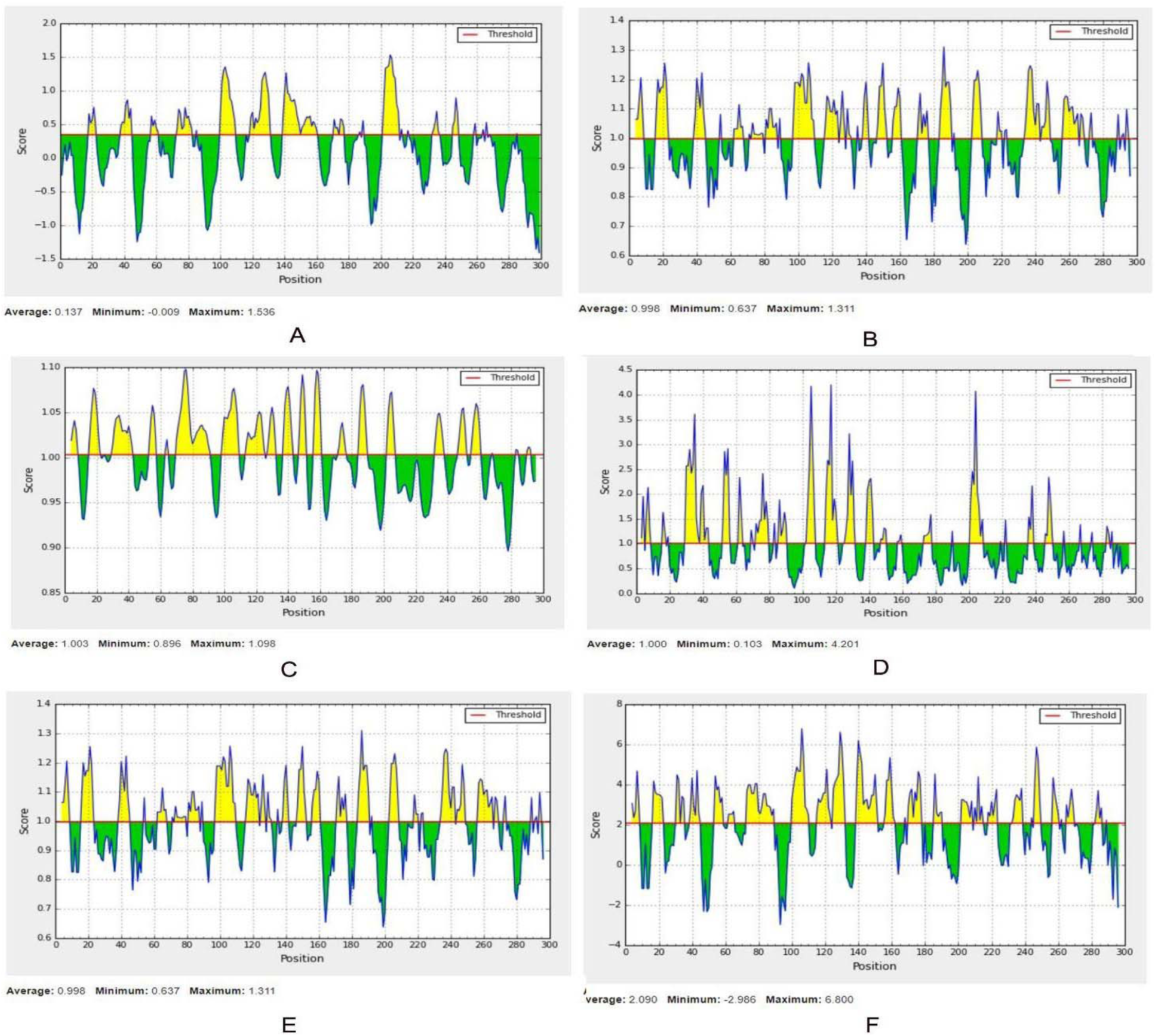
B-cell epitope prediction of Flagellar hook-associated protein (A: Linear, B: Beta-turn, C: Flexibility, D: Surface Accessibility, E: Antigenicity, F: Hydrophilicity). For each graph, X-axis and Y-axis represent the position and score. Residues that fall above the threshold value are shown in yellow color while the highest peak in yellow color identifies most favored position.

**Supplementary Figure 3.**
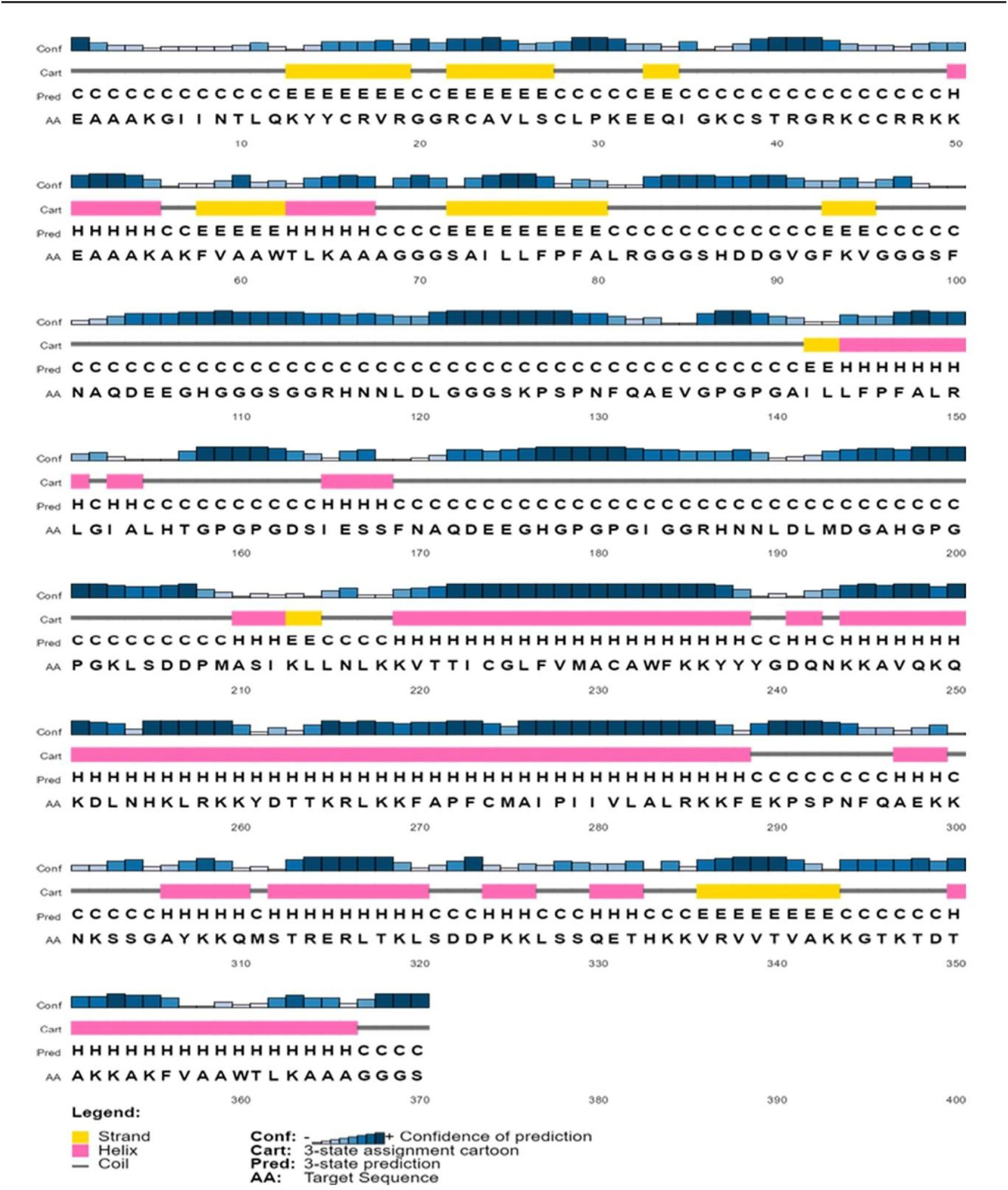
Secondary structure prediction of constructed vaccine protein V1 using PESIPRED server.

**Supplementary Figure 4.**
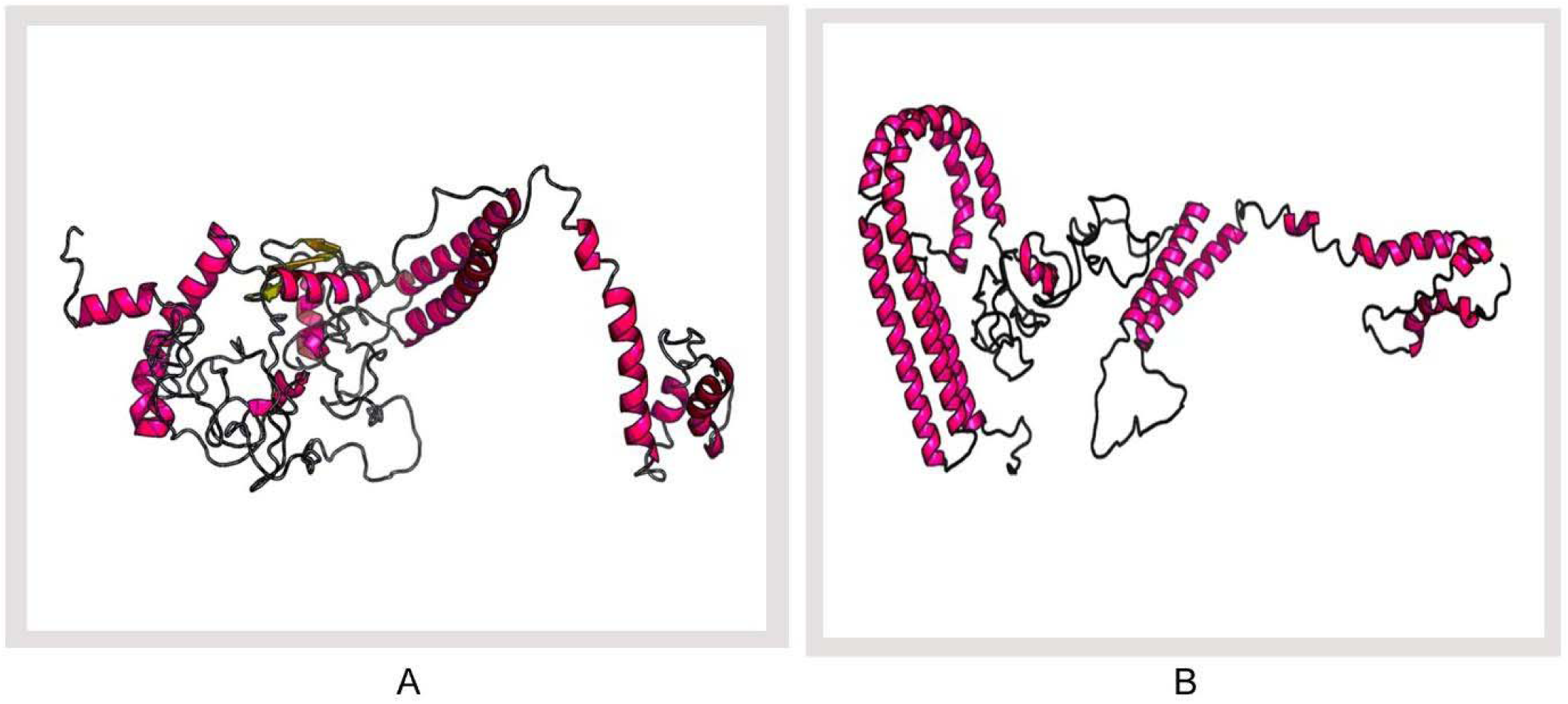
3D modeled structure of vaccine protein. (A) V2 and (B) V3 generated via RaptorX server.

**Supplementary Figure 5.**
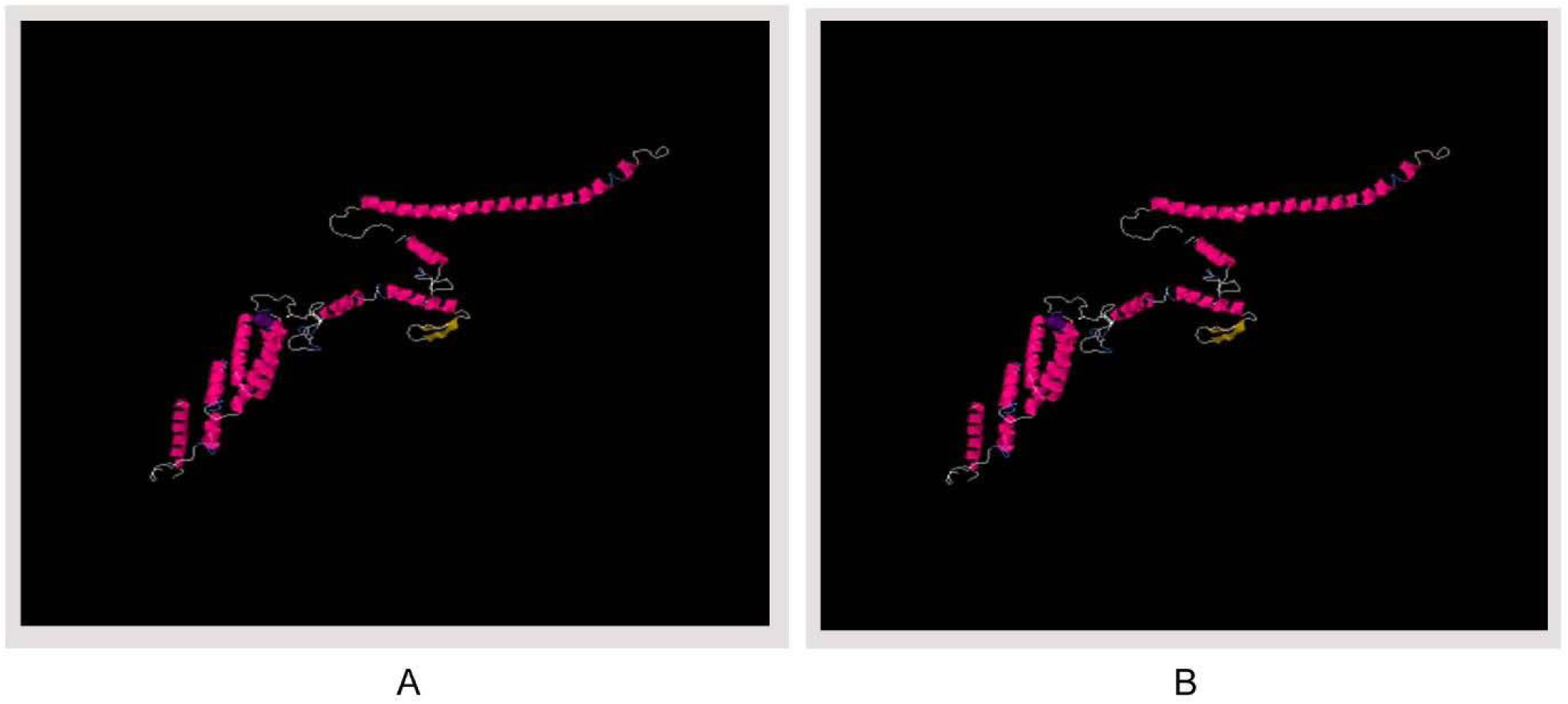
Disulfide engineering of vaccine protein V1. (A) Initial form, (B) Mutant form.

## Supplementary Tables

**Supplementary Table 1:**
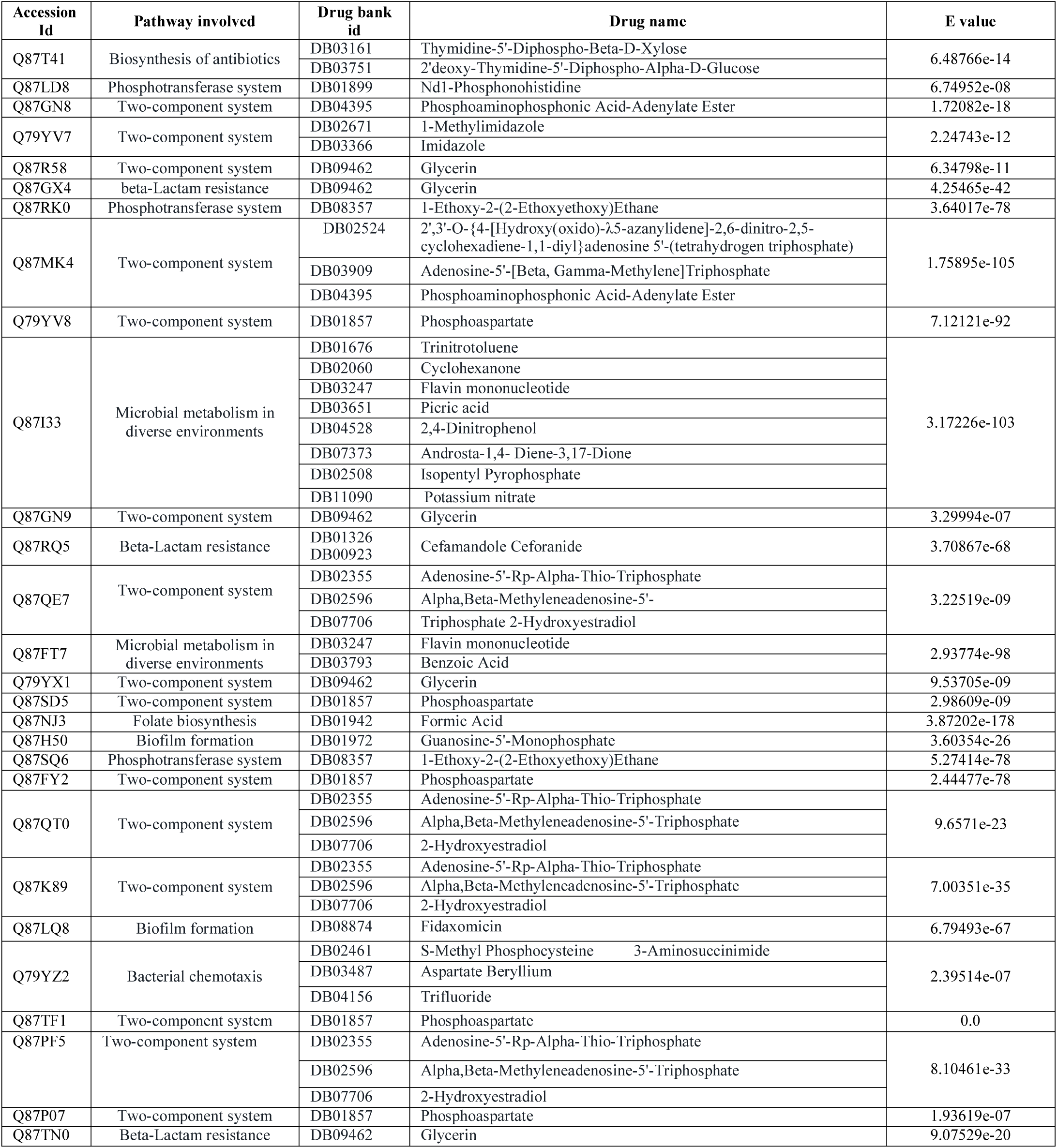
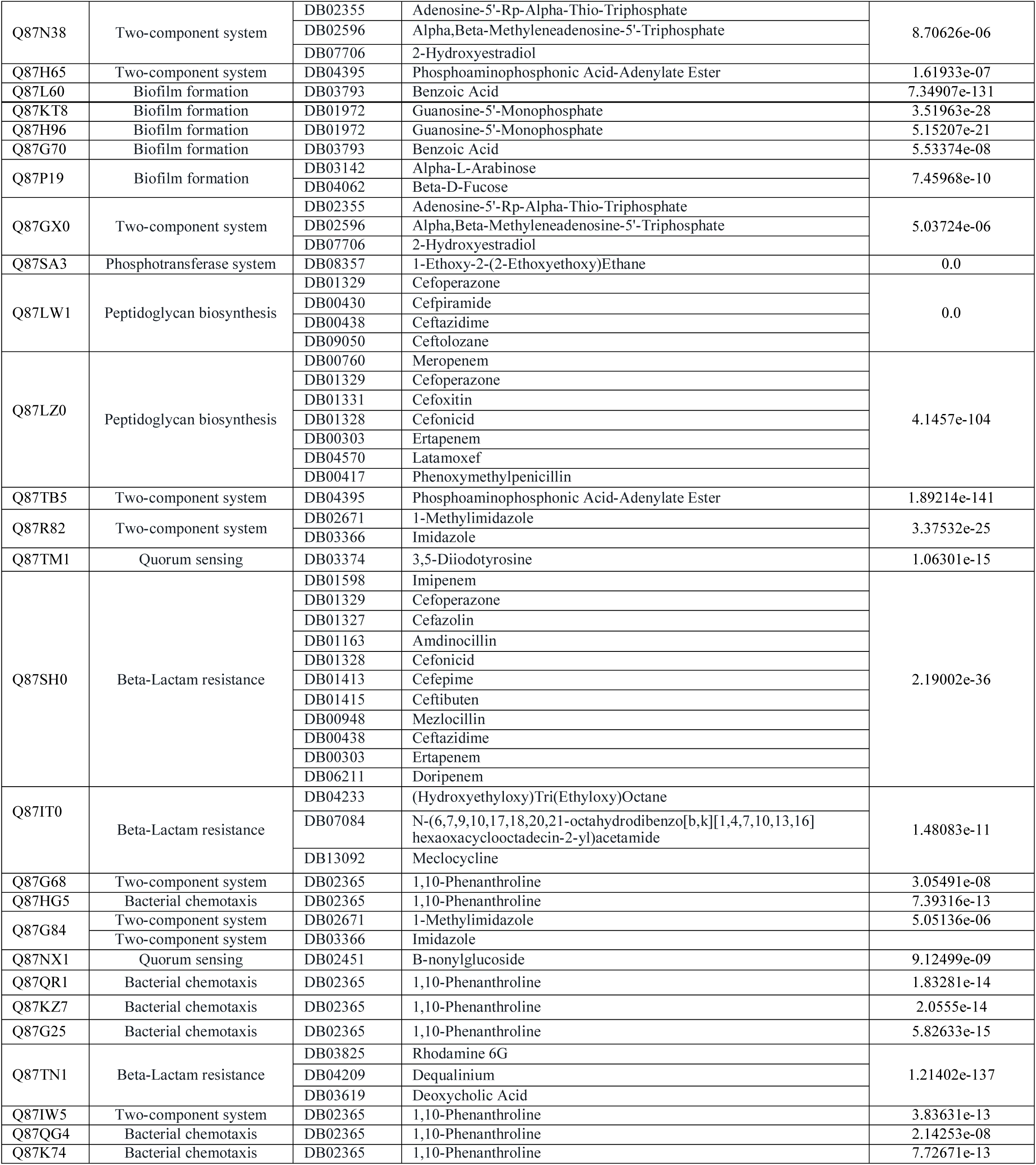
Pathway dependent metabolic proteins with druggable properties.

**Supplementary Table 2:**
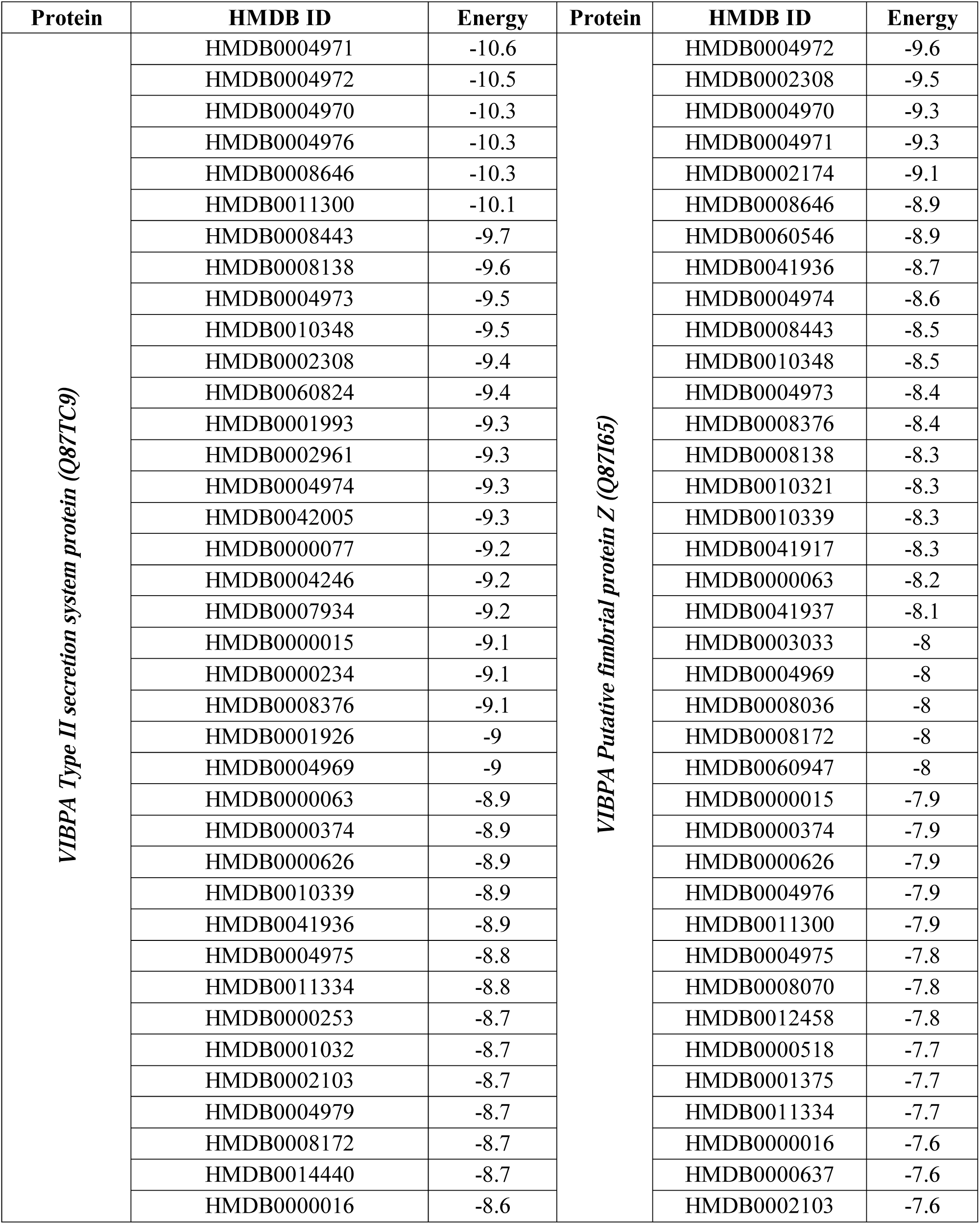
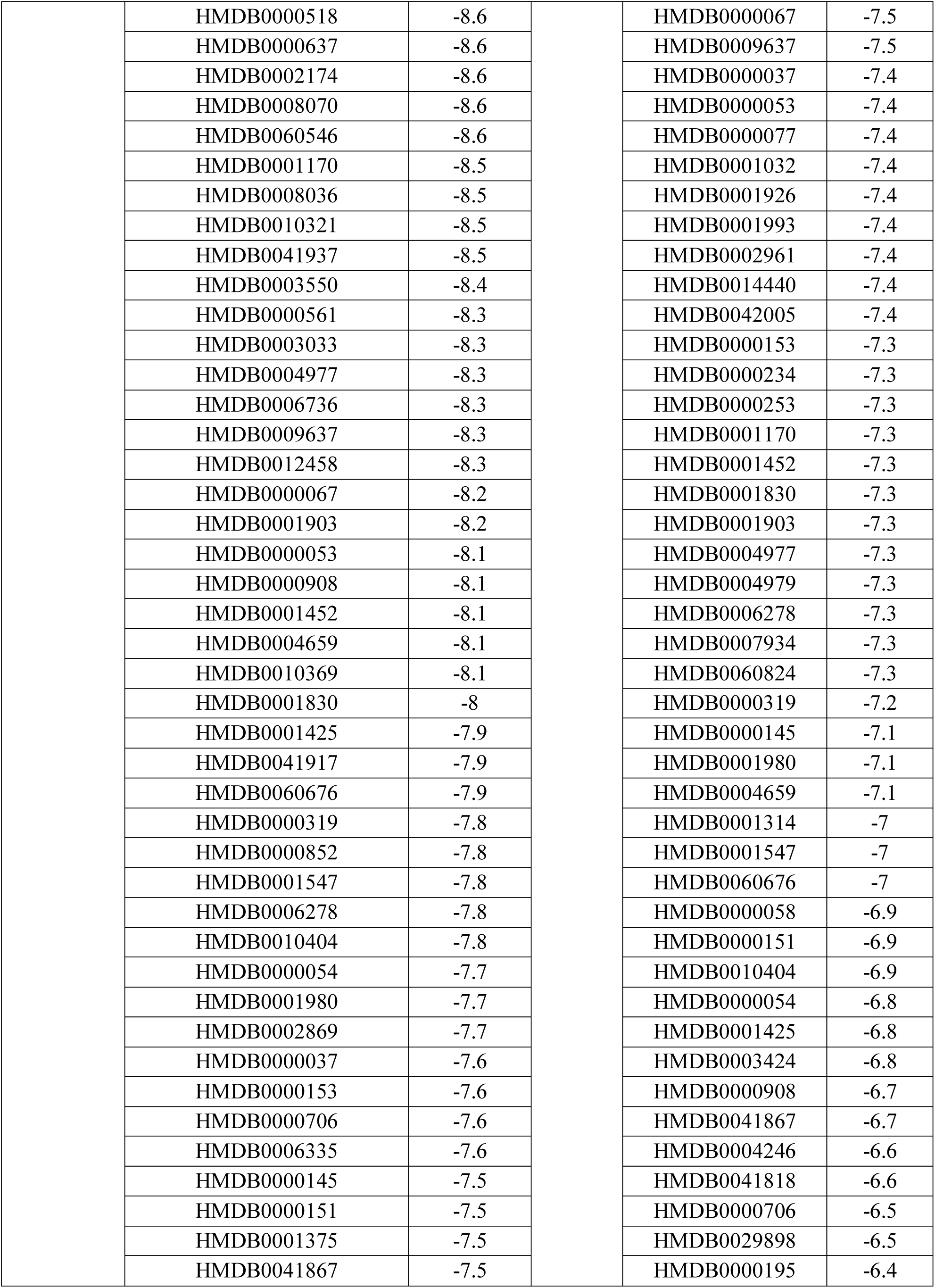
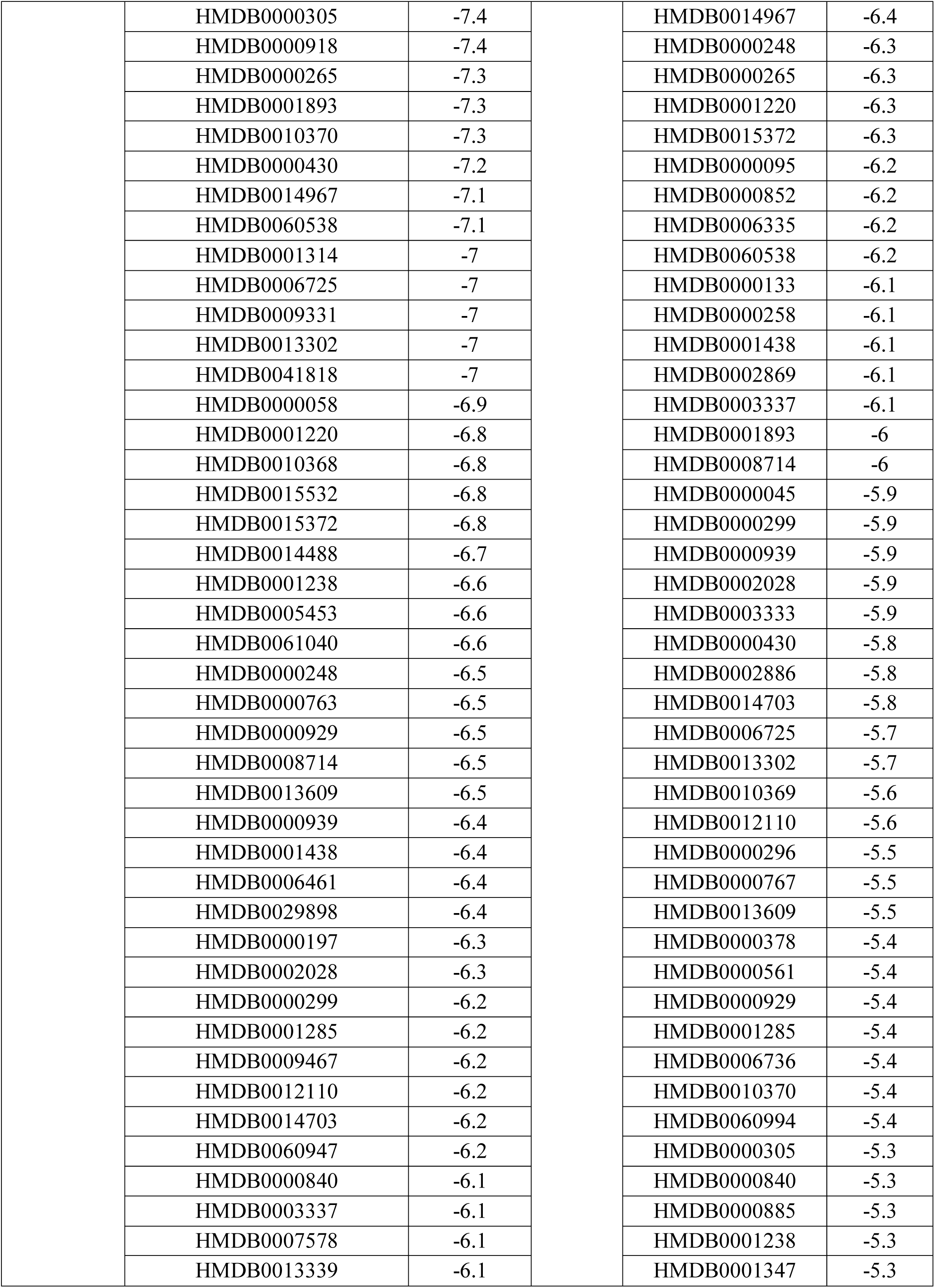
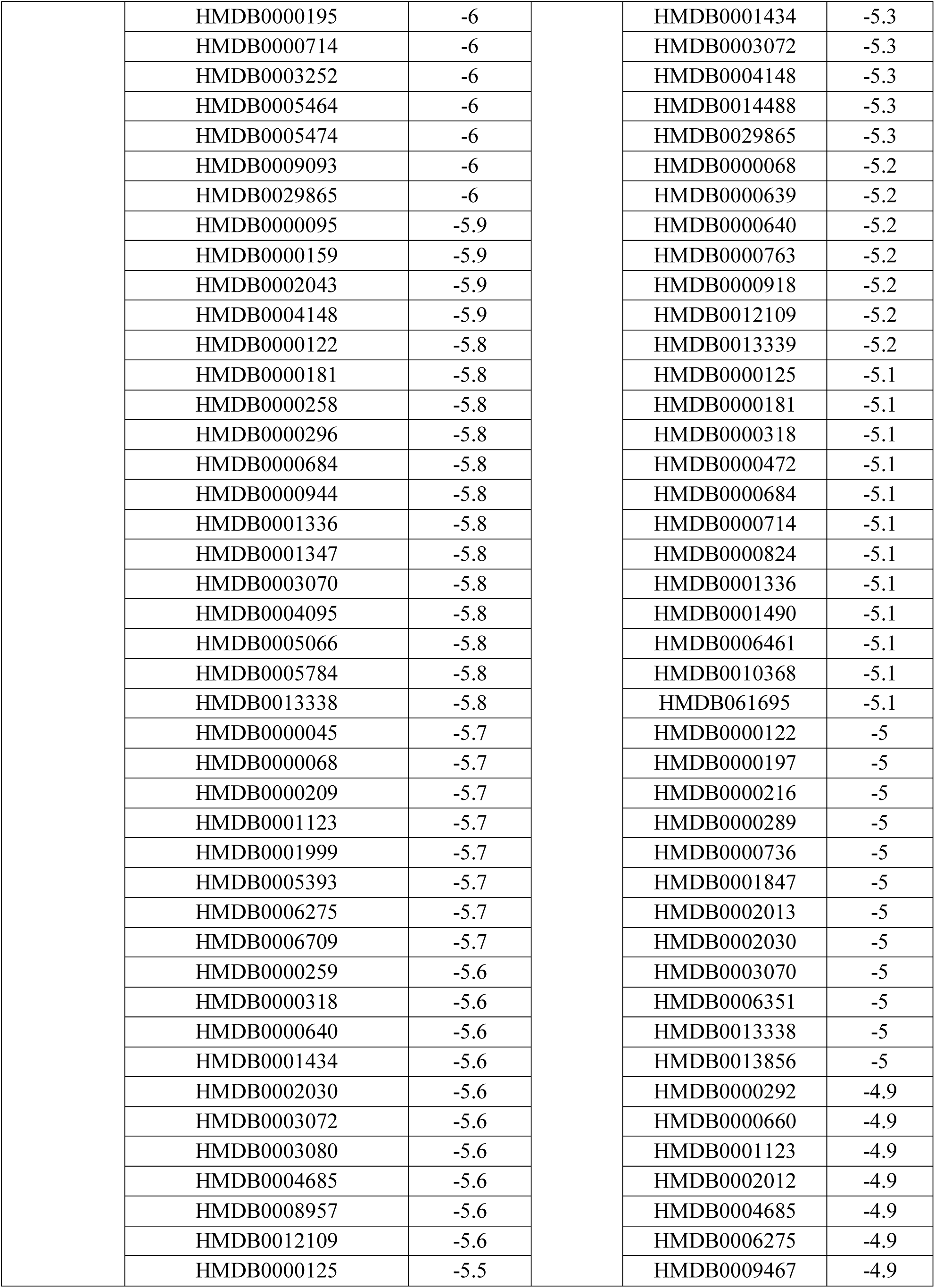
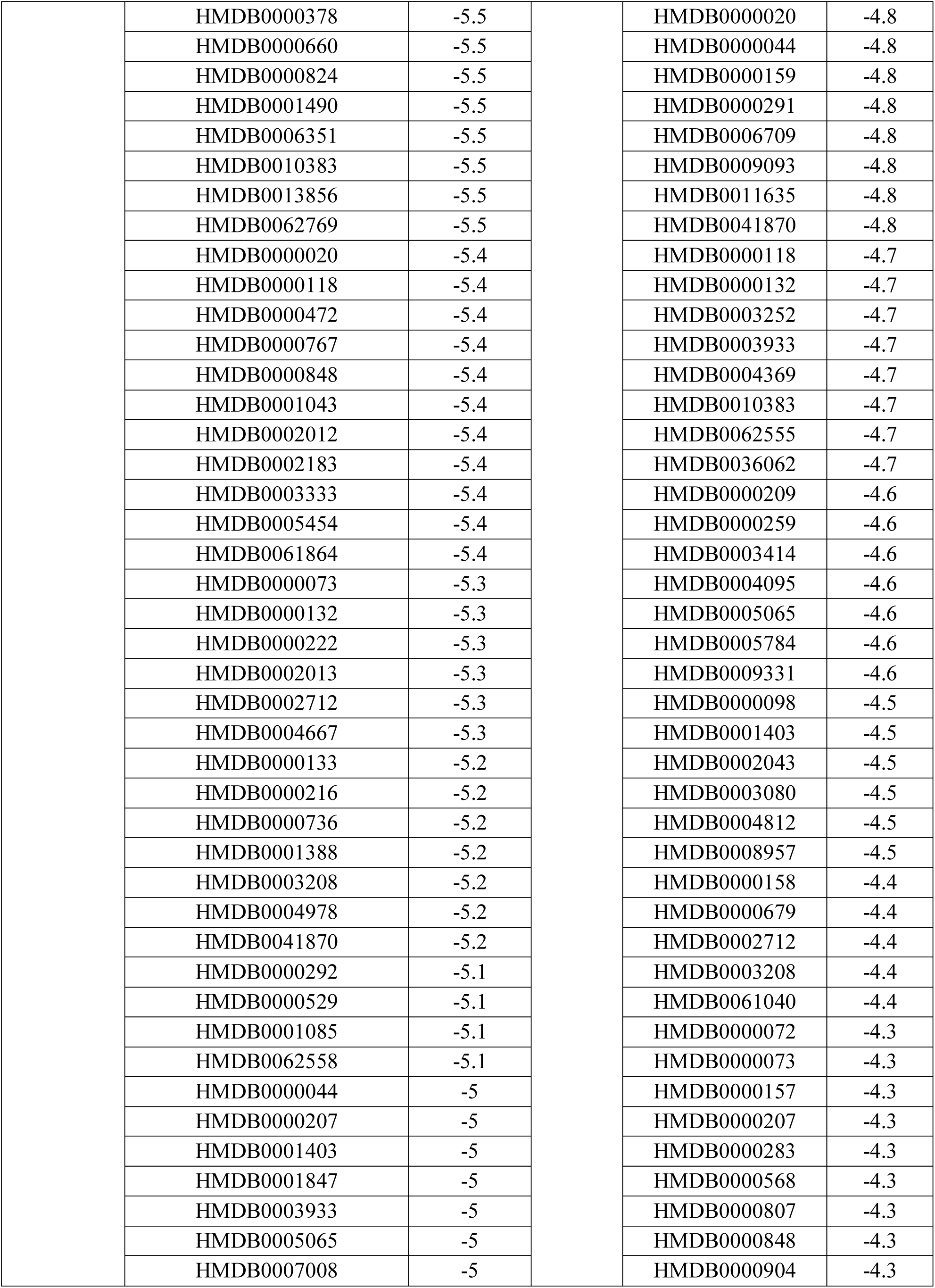
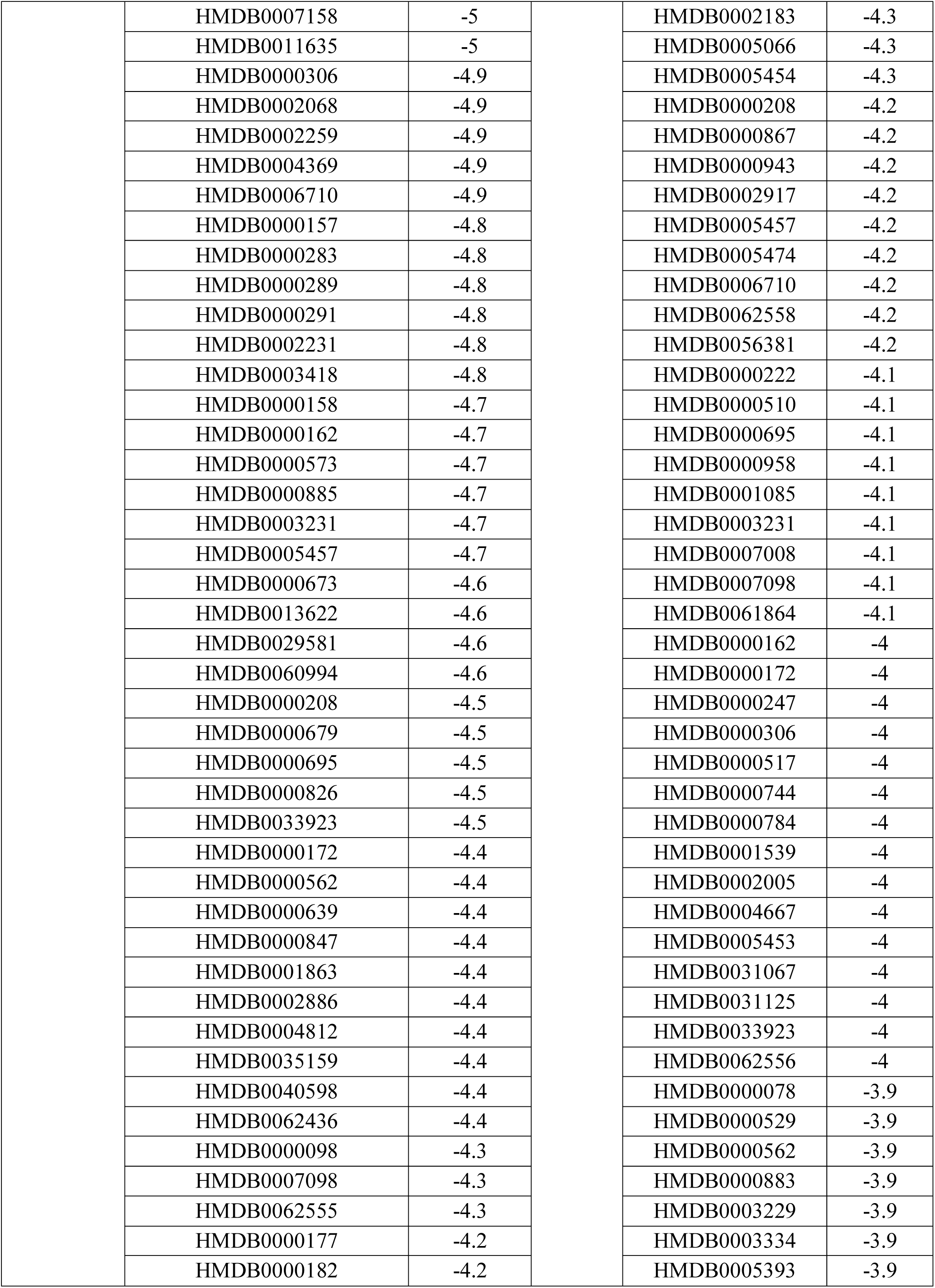
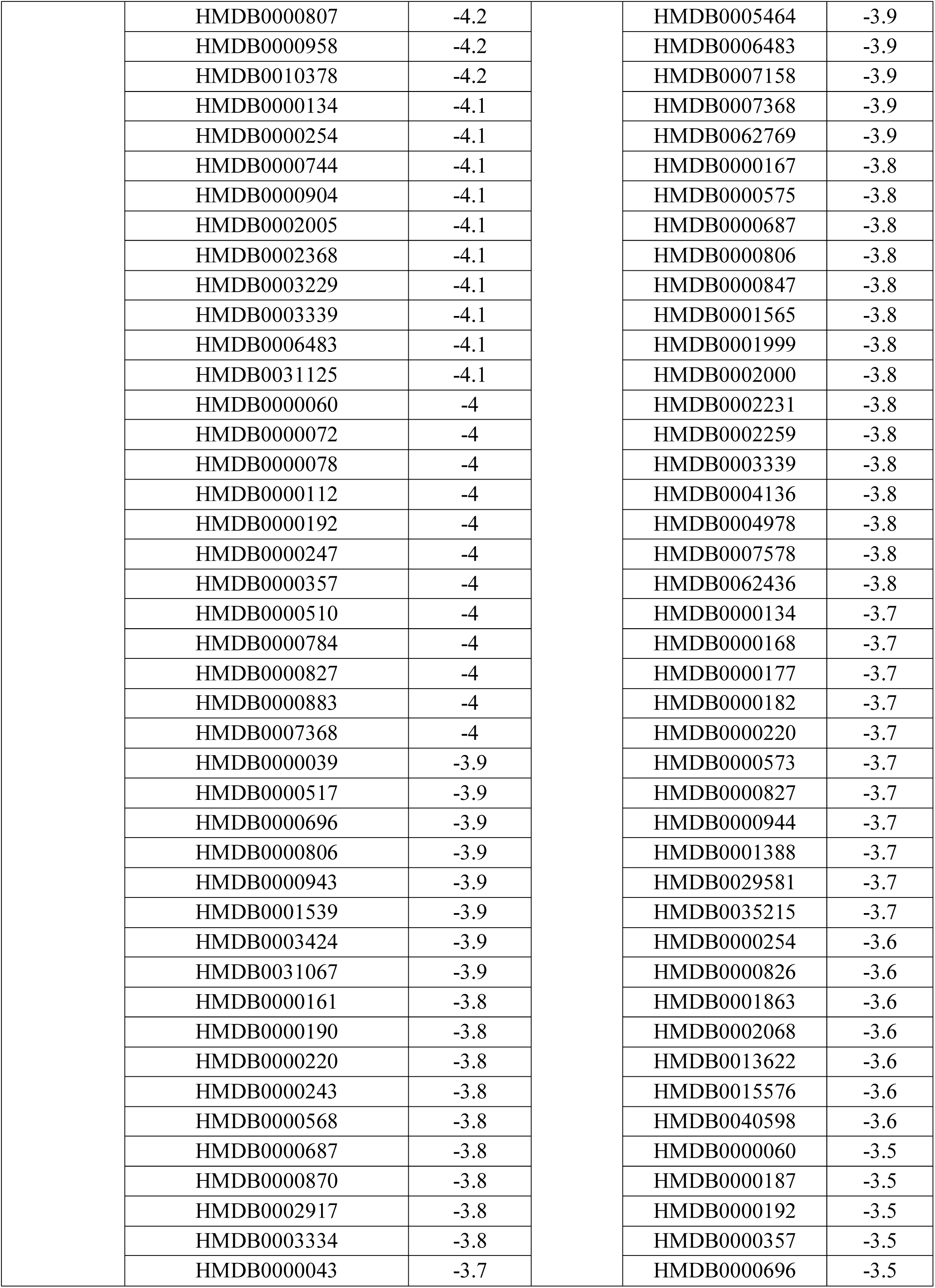
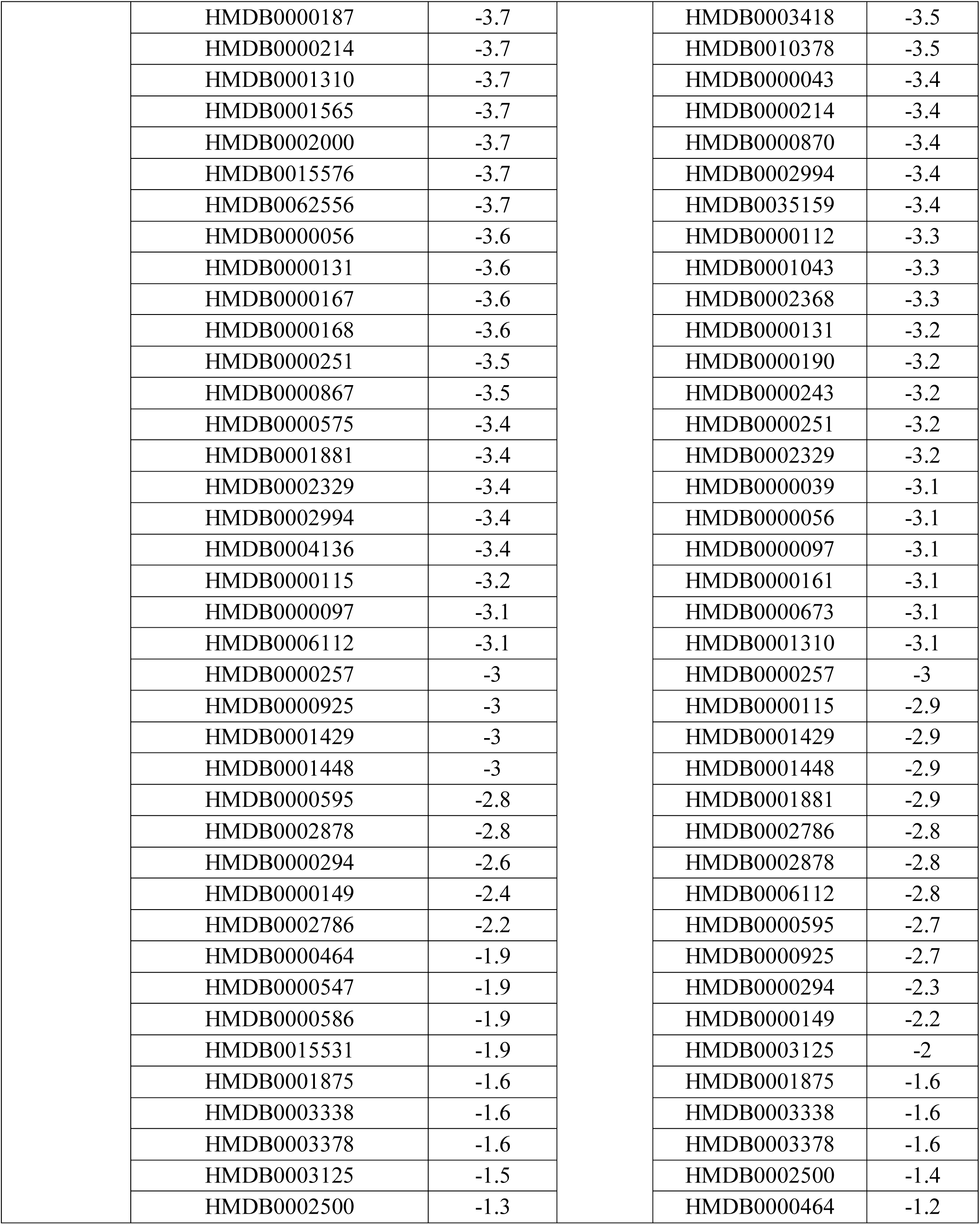
Predicted binding energy (docking score) of novel cytoplasmic proteins with human metabolites

| Protein | HMDB ID | Energy | Protein | HMDB ID | Energy |
| --- | --- | --- | --- | --- | --- |
| *VIBPA Type II secretion system protein (Q87TC9)* | HMDB0004971 | -10.6 | *VIBPA Putative fimbrial protein Z (Q87I65)* | HMDB0004972 | -9.6 |
|  | HMDB0004972 | -10.5 |  | HMDB0002308 | -9.5 |
|  | HMDB0004970 | -10.3 |  | HMDB0004970 | -9.3 |
|  | HMDB0004976 | -10.3 |  | HMDB0004971 | -9.3 |
|  | HMDB0008646 | -10.3 |  | HMDB0002174 | -9.1 |
|  | HMDB0011300 | -10.1 |  | HMDB0008646 | -8.9 |
|  | HMDB0008443 | -9.7 |  | HMDB0060546 | -8.9 |
|  | HMDB0008138 | -9.6 |  | HMDB0041936 | -8.7 |
|  | HMDB0004973 | -9.5 |  | HMDB0004974 | -8.6 |
|  | HMDB0010348 | -9.5 |  | HMDB0008443 | -8.5 |
|  | HMDB0002308 | -9.4 |  | HMDB0010348 | -8.5 |
|  | HMDB0060824 | -9.4 |  | HMDB0004973 | -8.4 |
|  | HMDB0001993 | -9.3 |  | HMDB0008376 | -8.4 |
|  | HMDB0002961 | -9.3 |  | HMDB0008138 | -8.3 |
|  | HMDB0004974 | -9.3 |  | HMDB0010321 | -8.3 |
|  | HMDB0042005 | -9.3 |  | HMDB0010339 | -8.3 |
|  | HMDB0000077 | -9.2 |  | HMDB0041917 | -8.3 |
|  | HMDB0004246 | -9.2 |  | HMDB0000063 | -8.2 |
|  | HMDB0007934 | -9.2 |  | HMDB0041937 | -8.1 |
|  | HMDB0000015 | -9.1 |  | HMDB0003033 | -8 |
|  | HMDB0000234 | -9.1 |  | HMDB0004969 | -8 |
|  | HMDB0008376 | -9.1 |  | HMDB0008036 | -8 |
|  | HMDB0001926 | -9 |  | HMDB0008172 | -8 |
|  | HMDB0004969 | -9 |  | HMDB0060947 | -8 |
|  | HMDB0000063 | -8.9 |  | HMDB0000015 | -7.9 |
|  | HMDB0000374 | -8.9 |  | HMDB0000374 | -7.9 |
|  | HMDB0000626 | -8.9 |  | HMDB0000626 | -7.9 |
|  | HMDB0010339 | -8.9 |  | HMDB0004976 | -7.9 |
|  | HMDB0041936 | -8.9 |  | HMDB0011300 | -7.9 |
|  | HMDB0004975 | -8.8 |  | HMDB0004975 | -7.8 |
|  | HMDB0011334 | -8.8 |  | HMDB0008070 | -7.8 |
|  | HMDB0000253 | -8.7 |  | HMDB0012458 | -7.8 |
|  | HMDB0001032 | -8.7 |  | HMDB0000518 | -7.7 |
|  | HMDB0002103 | -8.7 |  | HMDB0001375 | -7.7 |
|  | HMDB0004979 | -8.7 |  | HMDB0011334 | -7.7 |
|  | HMDB0008172 | -8.7 |  | HMDB0000016 | -7.6 |
|  | HMDB0014440 | -8.7 |  | HMDB0000637 | -7.6 |
|  | HMDB0000016 | -8.6 |  | HMDB0002103 | -7.6 |
|  | HMDB0000518 | -8.6 |  | HMDB0000067 | -7.5 |
|  | HMDB0000637 | -8.6 |  | HMDB0009637 | -7.5 |
|  | HMDB0002174 | -8.6 |  | HMDB0000037 | -7.4 |
|  | HMDB0008070 | -8.6 |  | HMDB0000053 | -7.4 |
|  | HMDB0060546 | -8.6 |  | HMDB0000077 | -7.4 |
|  | HMDB0001170 | -8.5 |  | HMDB0001032 | -7.4 |
|  | HMDB0008036 | -8.5 |  | HMDB0001926 | -7.4 |
|  | HMDB0010321 | -8.5 |  | HMDB0001993 | -7.4 |
|  | HMDB0041937 | -8.5 |  | HMDB0002961 | -7.4 |
|  | HMDB0003550 | -8.4 |  | HMDB0014440 | -7.4 |
|  | HMDB0000561 | -8.3 |  | HMDB0042005 | -7.4 |
|  | HMDB0003033 | -8.3 |  | HMDB0000153 | -7.3 |
|  | HMDB0004977 | -8.3 |  | HMDB0000234 | -7.3 |
|  | HMDB0006736 | -8.3 |  | HMDB0000253 | -7.3 |
|  | HMDB0009637 | -8.3 |  | HMDB0001170 | -7.3 |
|  | HMDB0012458 | -8.3 |  | HMDB0001452 | -7.3 |
|  | HMDB0000067 | -8.2 |  | HMDB0001830 | -7.3 |
|  | HMDB0001903 | -8.2 |  | HMDB0001903 | -7.3 |
|  | HMDB0000053 | -8.1 |  | HMDB0004977 | -7.3 |
|  | HMDB0000908 | -8.1 |  | HMDB0004979 | -7.3 |
|  | HMDB0001452 | -8.1 |  | HMDB0006278 | -7.3 |
|  | HMDB0004659 | -8.1 |  | HMDB0007934 | -7.3 |
|  | HMDB0010369 | -8.1 |  | HMDB0060824 | -7.3 |
|  | HMDB0001830 | -8 |  | HMDB0000319 | -7.2 |
|  | HMDB0001425 | -7.9 |  | HMDB0000145 | -7.1 |
|  | HMDB0041917 | -7.9 |  | HMDB0001980 | -7.1 |
|  | HMDB0060676 | -7.9 |  | HMDB0004659 | -7.1 |
|  | HMDB0000319 | -7.8 |  | HMDB0001314 | -7 |
|  | HMDB0000852 | -7.8 |  | HMDB0001547 | -7 |
|  | HMDB0001547 | -7.8 |  | HMDB0060676 | -7 |
|  | HMDB0006278 | -7.8 |  | HMDB0000058 | -6.9 |
|  | HMDB0010404 | -7.8 |  | HMDB0000151 | -6.9 |
|  | HMDB0000054 | -7.7 |  | HMDB0010404 | -6.9 |
|  | HMDB0001980 | -7.7 |  | HMDB0000054 | -6.8 |
|  | HMDB0002869 | -7.7 |  | HMDB0001425 | -6.8 |
|  | HMDB0000037 | -7.6 |  | HMDB0003424 | -6.8 |
|  | HMDB0000153 | -7.6 |  | HMDB0000908 | -6.7 |
|  | HMDB0000706 | -7.6 |  | HMDB0041867 | -6.7 |
|  | HMDB0006335 | -7.6 |  | HMDB0004246 | -6.6 |
|  | HMDB0000145 | -7.5 |  | HMDB0041818 | -6.6 |
|  | HMDB0000151 | -7.5 |  | HMDB0000706 | -6.5 |
|  | HMDB0001375 | -7.5 |  | HMDB0029898 | -6.5 |
|  | HMDB0041867 | -7.5 |  | HMDB0000195 | -6.4 |
|  | HMDB0000305 | -7.4 |  | HMDB0014967 | -6.4 |
|  | HMDB0000918 | -7.4 |  | HMDB0000248 | -6.3 |
|  | HMDB0000265 | -7.3 |  | HMDB0000265 | -6.3 |
|  | HMDB0001893 | -7.3 |  | HMDB0001220 | -6.3 |
|  | HMDB0010370 | -7.3 |  | HMDB0015372 | -6.3 |
|  | HMDB0000430 | -7.2 |  | HMDB0000095 | -6.2 |
|  | HMDB0014967 | -7.1 |  | HMDB0000852 | -6.2 |
|  | HMDB0060538 | -7.1 |  | HMDB0006335 | -6.2 |
|  | HMDB0001314 | -7 |  | HMDB0060538 | -6.2 |
|  | HMDB0006725 | -7 |  | HMDB0000133 | -6.1 |
|  | HMDB0009331 | -7 |  | HMDB0000258 | -6.1 |
|  | HMDB0013302 | -7 |  | HMDB0001438 | -6.1 |
|  | HMDB0041818 | -7 |  | HMDB0002869 | -6.1 |
|  | HMDB0000058 | -6.9 |  | HMDB0003337 | -6.1 |
|  | HMDB0001220 | -6.8 |  | HMDB0001893 | -6 |
|  | HMDB0010368 | -6.8 |  | HMDB0008714 | -6 |
|  | HMDB0015532 | -6.8 |  | HMDB0000045 | -5.9 |
|  | HMDB0015372 | -6.8 |  | HMDB0000299 | -5.9 |
|  | HMDB0014488 | -6.7 |  | HMDB0000939 | -5.9 |
|  | HMDB0001238 | -6.6 |  | HMDB0002028 | -5.9 |
|  | HMDB0005453 | -6.6 |  | HMDB0003333 | -5.9 |
|  | HMDB0061040 | -6.6 |  | HMDB0000430 | -5.8 |
|  | HMDB0000248 | -6.5 |  | HMDB0002886 | -5.8 |
|  | HMDB0000763 | -6.5 |  | HMDB0014703 | -5.8 |
|  | HMDB0000929 | -6.5 |  | HMDB0006725 | -5.7 |
|  | HMDB0008714 | -6.5 |  | HMDB0013302 | -5.7 |
|  | HMDB0013609 | -6.5 |  | HMDB0010369 | -5.6 |
|  | HMDB0000939 | -6.4 |  | HMDB0012110 | -5.6 |
|  | HMDB0001438 | -6.4 |  | HMDB0000296 | -5.5 |
|  | HMDB0006461 | -6.4 |  | HMDB0000767 | -5.5 |
|  | HMDB0029898 | -6.4 |  | HMDB0013609 | -5.5 |
|  | HMDB0000197 | -6.3 |  | HMDB0000378 | -5.4 |
|  | HMDB0002028 | -6.3 |  | HMDB0000561 | -5.4 |
|  | HMDB0000299 | -6.2 |  | HMDB0000929 | -5.4 |
|  | HMDB0001285 | -6.2 |  | HMDB0001285 | -5.4 |
|  | HMDB0009467 | -6.2 |  | HMDB0006736 | -5.4 |
|  | HMDB0012110 | -6.2 |  | HMDB0010370 | -5.4 |
|  | HMDB0014703 | -6.2 |  | HMDB0060994 | -5.4 |
|  | HMDB0060947 | -6.2 |  | HMDB0000305 | -5.3 |
|  | HMDB0000840 | -6.1 |  | HMDB0000840 | -5.3 |
|  | HMDB0003337 | -6.1 |  | HMDB0000885 | -5.3 |
|  | HMDB0007578 | -6.1 |  | HMDB0001238 | -5.3 |
|  | HMDB0013339 | -6.1 |  | HMDB0001347 | -5.3 |

|  |  |  |  |
| --- | --- | --- | --- |
| HMDB0000195 | -6 | HMDB0001434 | -5.3 |
| HMDB0000714 | -6 | HMDB0003072 | -5.3 |
| HMDB0003252 | -6 | HMDB0004148 | -5.3 |
| HMDB0005464 | -6 | HMDB0014488 | -5.3 |
| HMDB0005474 | -6 | HMDB0029865 | -5.3 |
| HMDB0009093 | -6 | HMDB0000068 | -5.2 |
| HMDB0029865 | -6 | HMDB0000639 | -5.2 |
| HMDB0000095 | -5.9 | HMDB0000640 | -5.2 |
| HMDB0000159 | -5.9 | HMDB0000763 | -5.2 |
| HMDB0002043 | -5.9 | HMDB0000918 | -5.2 |
| HMDB0004148 | -5.9 | HMDB0012109 | -5.2 |
| HMDB0000122 | -5.8 | HMDB0013339 | -5.2 |
| HMDB0000181 | -5.8 | HMDB0000125 | -5.1 |
| HMDB0000258 | -5.8 | HMDB0000181 | -5.1 |
| HMDB0000296 | -5.8 | HMDB0000318 | -5.1 |
| HMDB0000684 | -5.8 | HMDB0000472 | -5.1 |
| HMDB0000944 | -5.8 | HMDB0000684 | -5.1 |
| HMDB0001336 | -5.8 | HMDB0000714 | -5.1 |
| HMDB0001347 | -5.8 | HMDB0000824 | -5.1 |
| HMDB0003070 | -5.8 | HMDB0001336 | -5.1 |
| HMDB0004095 | -5.8 | HMDB0001490 | -5.1 |
| HMDB0005066 | -5.8 | HMDB0006461 | -5.1 |
| HMDB0005784 | -5.8 | HMDB0010368 | -5.1 |
| HMDB0013338 | -5.8 | HMDB061695 | -5.1 |
| HMDB0000045 | -5.7 | HMDB0000122 | -5 |
| HMDB0000068 | -5.7 | HMDB0000197 | -5 |
| HMDB0000209 | -5.7 | HMDB0000216 | -5 |
| HMDB0001123 | -5.7 | HMDB0000289 | -5 |
| HMDB0001999 | -5.7 | HMDB0000736 | -5 |
| HMDB0005393 | -5.7 | HMDB0001847 | -5 |
| HMDB0006275 | -5.7 | HMDB0002013 | -5 |
| HMDB0006709 | -5.7 | HMDB0002030 | -5 |
| HMDB0000259 | -5.6 | HMDB0003070 | -5 |
| HMDB0000318 | -5.6 | HMDB0006351 | -5 |
| HMDB0000640 | -5.6 | HMDB0013338 | -5 |
| HMDB0001434 | -5.6 | HMDB0013856 | -5 |
| HMDB0002030 | -5.6 | HMDB0000292 | -4.9 |
| HMDB0003072 | -5.6 | HMDB0000660 | -4.9 |
| HMDB0003080 | -5.6 | HMDB0001123 | -4.9 |
| HMDB0004685 | -5.6 | HMDB0002012 | -4.9 |
| HMDB0008957 | -5.6 | HMDB0004685 | -4.9 |
| HMDB0012109 | -5.6 | HMDB0006275 | -4.9 |
| HMDB0000125 | -5.5 | HMDB0009467 | -4.9 |
| HMDB0000378 | -5.5 | HMDB0000020 | -4.8 |
| HMDB0000660 | -5.5 | HMDB0000044 | -4.8 |
| HMDB0000824 | -5.5 | HMDB0000159 | -4.8 |
| HMDB0001490 | -5.5 | HMDB0000291 | -4.8 |
| HMDB0006351 | -5.5 | HMDB0006709 | -4.8 |
| HMDB0010383 | -5.5 | HMDB0009093 | -4.8 |
| HMDB0013856 | -5.5 | HMDB0011635 | -4.8 |
| HMDB0062769 | -5.5 | HMDB0041870 | -4.8 |
| HMDB0000020 | -5.4 | HMDB0000118 | -4.7 |
| HMDB0000118 | -5.4 | HMDB0000132 | -4.7 |
| HMDB0000472 | -5.4 | HMDB0003252 | -4.7 |
| HMDB0000767 | -5.4 | HMDB0003933 | -4.7 |
| HMDB0000848 | -5.4 | HMDB0004369 | -4.7 |
| HMDB0001043 | -5.4 | HMDB0010383 | -4.7 |
| HMDB0002012 | -5.4 | HMDB0062555 | -4.7 |
| HMDB0002183 | -5.4 | HMDB0036062 | -4.7 |
| HMDB0003333 | -5.4 | HMDB0000209 | -4.6 |
| HMDB0005454 | -5.4 | HMDB0000259 | -4.6 |
| HMDB0061864 | -5.4 | HMDB0003414 | -4.6 |
| HMDB0000073 | -5.3 | HMDB0004095 | -4.6 |
| HMDB0000132 | -5.3 | HMDB0005065 | -4.6 |
| HMDB0000222 | -5.3 | HMDB0005784 | -4.6 |
| HMDB0002013 | -5.3 | HMDB0009331 | -4.6 |
| HMDB0002712 | -5.3 | HMDB0000098 | -4.5 |
| HMDB0004667 | -5.3 | HMDB0001403 | -4.5 |
| HMDB0000133 | -5.2 | HMDB0002043 | -4.5 |
| HMDB0000216 | -5.2 | HMDB0003080 | -4.5 |
| HMDB0000736 | -5.2 | HMDB0004812 | -4.5 |
| HMDB0001388 | -5.2 | HMDB0008957 | -4.5 |
| HMDB0003208 | -5.2 | HMDB0000158 | -4.4 |
| HMDB0004978 | -5.2 | HMDB0000679 | -4.4 |
| HMDB0041870 | -5.2 | HMDB0002712 | -4.4 |
| HMDB0000292 | -5.1 | HMDB0003208 | -4.4 |
| HMDB0000529 | -5.1 | HMDB0061040 | -4.4 |
| HMDB0001085 | -5.1 | HMDB0000072 | -4.3 |
| HMDB0062558 | -5.1 | HMDB0000073 | -4.3 |
| HMDB0000044 | -5 | HMDB0000157 | -4.3 |
| HMDB0000207 | -5 | HMDB0000207 | -4.3 |
| HMDB0001403 | -5 | HMDB0000283 | -4.3 |
| HMDB0001847 | -5 | HMDB0000568 | -4.3 |
| HMDB0003933 | -5 | HMDB0000807 | -4.3 |
| HMDB0005065 | -5 | HMDB0000848 | -4.3 |
| HMDB0007008 | -5 | HMDB0000904 | -4.3 |
|  | HMDB0007158 | -5 |  | HMDB0002183 | -4.3 |
|  | HMDB0011635 | -5 |  | HMDB0005066 | -4.3 |
|  | HMDB0000306 | -4.9 |  | HMDB0005454 | -4.3 |
|  | HMDB0002068 | -4.9 |  | HMDB0000208 | -4.2 |
|  | HMDB0002259 | -4.9 |  | HMDB0000867 | -4.2 |
|  | HMDB0004369 | -4.9 |  | HMDB0000943 | -4.2 |
|  | HMDB0006710 | -4.9 |  | HMDB0002917 | -4.2 |
|  | HMDB0000157 | -4.8 |  | HMDB0005457 | -4.2 |
|  | HMDB0000283 | -4.8 |  | HMDB0005474 | -4.2 |
|  | HMDB0000289 | -4.8 |  | HMDB0006710 | -4.2 |
|  | HMDB0000291 | -4.8 |  | HMDB0062558 | -4.2 |
|  | HMDB0002231 | -4.8 |  | HMDB0056381 | -4.2 |
|  | HMDB0003418 | -4.8 |  | HMDB0000222 | -4.1 |
|  | HMDB0000158 | -4.7 |  | HMDB0000510 | -4.1 |
|  | HMDB0000162 | -4.7 |  | HMDB0000695 | -4.1 |
|  | HMDB0000573 | -4.7 |  | HMDB0000958 | -4.1 |
|  | HMDB0000885 | -4.7 |  | HMDB0001085 | -4.1 |
|  | HMDB0003231 | -4.7 |  | HMDB0003231 | -4.1 |
|  | HMDB0005457 | -4.7 |  | HMDB0007008 | -4.1 |
|  | HMDB0000673 | -4.6 |  | HMDB0007098 | -4.1 |
|  | HMDB0013622 | -4.6 |  | HMDB0061864 | -4.1 |
|  | HMDB0029581 | -4.6 |  | HMDB0000162 | -4 |
|  | HMDB0060994 | -4.6 |  | HMDB0000172 | -4 |
|  | HMDB0000208 | -4.5 |  | HMDB0000247 | -4 |
|  | HMDB0000679 | -4.5 |  | HMDB0000306 | -4 |
|  | HMDB0000695 | -4.5 |  | HMDB0000517 | -4 |
|  | HMDB0000826 | -4.5 |  | HMDB0000744 | -4 |
|  | HMDB0033923 | -4.5 |  | HMDB0000784 | -4 |
|  | HMDB0000172 | -4.4 |  | HMDB0001539 | -4 |
|  | HMDB0000562 | -4.4 |  | HMDB0002005 | -4 |
|  | HMDB0000639 | -4.4 |  | HMDB0004667 | -4 |
|  | HMDB0000847 | -4.4 |  | HMDB0005453 | -4 |
|  | HMDB0001863 | -4.4 |  | HMDB0031067 | -4 |
|  | HMDB0002886 | -4.4 |  | HMDB0031125 | -4 |
|  | HMDB0004812 | -4.4 |  | HMDB0033923 | -4 |
|  | HMDB0035159 | -4.4 |  | HMDB0062556 | -4 |
|  | HMDB0040598 | -4.4 |  | HMDB0000078 | -3.9 |
|  | HMDB0062436 | -4.4 |  | HMDB0000529 | -3.9 |
|  | HMDB0000098 | -4.3 |  | HMDB0000562 | -3.9 |
|  | HMDB0007098 | -4.3 |  | HMDB0000883 | -3.9 |
|  | HMDB0062555 | -4.3 |  | HMDB0003229 | -3.9 |
|  | HMDB0000177 | -4.2 |  | HMDB0003334 | -3.9 |
|  | HMDB0000182 | -4.2 |  | HMDB0005393 | -3.9 |

|  |  |  |  |
| --- | --- | --- | --- |
| HMDB0000807 | -4.2 | HMDB0005464 | -3.9 |
| HMDB0000958 | -4.2 | HMDB0006483 | -3.9 |
| HMDB0010378 | -4.2 | HMDB0007158 | -3.9 |
| HMDB0000134 | -4.1 | HMDB0007368 | -3.9 |
| HMDB0000254 | -4.1 | HMDB0062769 | -3.9 |
| HMDB0000744 | -4.1 | HMDB0000167 | -3.8 |
| HMDB0000904 | -4.1 | HMDB0000575 | -3.8 |
| HMDB0002005 | -4.1 | HMDB0000687 | -3.8 |
| HMDB0002368 | -4.1 | HMDB0000806 | -3.8 |
| HMDB0003229 | -4.1 | HMDB0000847 | -3.8 |
| HMDB0003339 | -4.1 | HMDB0001565 | -3.8 |
| HMDB0006483 | -4.1 | HMDB0001999 | -3.8 |
| HMDB0031125 | -4.1 | HMDB0002000 | -3.8 |
| HMDB0000060 | -4 | HMDB0002231 | -3.8 |
| HMDB0000072 | -4 | HMDB0002259 | -3.8 |
| HMDB0000078 | -4 | HMDB0003339 | -3.8 |
| HMDB0000112 | -4 | HMDB0004136 | -3.8 |
| HMDB0000192 | -4 | HMDB0004978 | -3.8 |
| HMDB0000247 | -4 | HMDB0007578 | -3.8 |
| HMDB0000357 | -4 | HMDB0062436 | -3.8 |
| HMDB0000510 | -4 | HMDB0000134 | -3.7 |
| HMDB0000784 | -4 | HMDB0000168 | -3.7 |
| HMDB0000827 | -4 | HMDB0000177 | -3.7 |
| HMDB0000883 | -4 | HMDB0000182 | -3.7 |
| HMDB0007368 | -4 | HMDB0000220 | -3.7 |
| HMDB0000039 | -3.9 | HMDB0000573 | -3.7 |
| HMDB0000517 | -3.9 | HMDB0000827 | -3.7 |
| HMDB0000696 | -3.9 | HMDB0000944 | -3.7 |
| HMDB0000806 | -3.9 | HMDB0001388 | -3.7 |
| HMDB0000943 | -3.9 | HMDB0029581 | -3.7 |
| HMDB0001539 | -3.9 | HMDB0035215 | -3.7 |
| HMDB0003424 | -3.9 | HMDB0000254 | -3.6 |
| HMDB0031067 | -3.9 | HMDB0000826 | -3.6 |
| HMDB0000161 | -3.8 | HMDB0001863 | -3.6 |
| HMDB0000190 | -3.8 | HMDB0002068 | -3.6 |
| HMDB0000220 | -3.8 | HMDB0013622 | -3.6 |
| HMDB0000243 | -3.8 | HMDB0015576 | -3.6 |
| HMDB0000568 | -3.8 | HMDB0040598 | -3.6 |
| HMDB0000687 | -3.8 | HMDB0000060 | -3.5 |
| HMDB0000870 | -3.8 | HMDB0000187 | -3.5 |
| HMDB0002917 | -3.8 | HMDB0000192 | -3.5 |
| HMDB0003334 | -3.8 | HMDB0000357 | -3.5 |
| HMDB0000043 | -3.7 | HMDB0000696 | -3.5 |
|  | HMDB0000187 | -3.7 |  | HMDB0003418 | -3.5 |
|  | HMDB0000214 | -3.7 |  | HMDB0010378 | -3.5 |
|  | HMDB0001310 | -3.7 |  | HMDB0000043 | -3.4 |
|  | HMDB0001565 | -3.7 |  | HMDB0000214 | -3.4 |
|  | HMDB0002000 | -3.7 |  | HMDB0000870 | -3.4 |
|  | HMDB0015576 | -3.7 |  | HMDB0002994 | -3.4 |
|  | HMDB0062556 | -3.7 |  | HMDB0035159 | -3.4 |
|  | HMDB0000056 | -3.6 |  | HMDB0000112 | -3.3 |
|  | HMDB0000131 | -3.6 |  | HMDB0001043 | -3.3 |
|  | HMDB0000167 | -3.6 |  | HMDB0002368 | -3.3 |
|  | HMDB0000168 | -3.6 |  | HMDB0000131 | -3.2 |
|  | HMDB0000251 | -3.5 |  | HMDB0000190 | -3.2 |
|  | HMDB0000867 | -3.5 |  | HMDB0000243 | -3.2 |
|  | HMDB0000575 | -3.4 |  | HMDB0000251 | -3.2 |
|  | HMDB0001881 | -3.4 |  | HMDB0002329 | -3.2 |
|  | HMDB0002329 | -3.4 |  | HMDB0000039 | -3.1 |
|  | HMDB0002994 | -3.4 |  | HMDB0000056 | -3.1 |
|  | HMDB0004136 | -3.4 |  | HMDB0000097 | -3.1 |
|  | HMDB0000115 | -3.2 |  | HMDB0000161 | -3.1 |
|  | HMDB0000097 | -3.1 |  | HMDB0000673 | -3.1 |
|  | HMDB0006112 | -3.1 |  | HMDB0001310 | -3.1 |
|  | HMDB0000257 | -3 |  | HMDB0000257 | -3 |
|  | HMDB0000925 | -3 |  | HMDB0000115 | -2.9 |
|  | HMDB0001429 | -3 |  | HMDB0001429 | -2.9 |
|  | HMDB0001448 | -3 |  | HMDB0001448 | -2.9 |
|  | HMDB0000595 | -2.8 |  | HMDB0001881 | -2.9 |
|  | HMDB0002878 | -2.8 |  | HMDB0002786 | -2.8 |
|  | HMDB0000294 | -2.6 |  | HMDB0002878 | -2.8 |
|  | HMDB0000149 | -2.4 |  | HMDB0006112 | -2.8 |
|  | HMDB0002786 | -2.2 |  | HMDB0000595 | -2.7 |
|  | HMDB0000464 | -1.9 |  | HMDB0000925 | -2.7 |
|  | HMDB0000547 | -1.9 |  | HMDB0000294 | -2.3 |
|  | HMDB0000586 | -1.9 |  | HMDB0000149 | -2.2 |
|  | HMDB0015531 | -1.9 |  | HMDB0003125 | -2 |
|  | HMDB0001875 | -1.6 |  | HMDB0001875 | -1.6 |
|  | HMDB0003338 | -1.6 |  | HMDB0003338 | -1.6 |
|  | HMDB0003378 | -1.6 |  | HMDB0003378 | -1.6 |
|  | HMDB0003125 | -1.5 |  | HMDB0002500 | -1.4 |
|  | HMDB0002500 | -1.3 |  | HMDB0000464 | -1.2 |

**Supplementary table 3:**
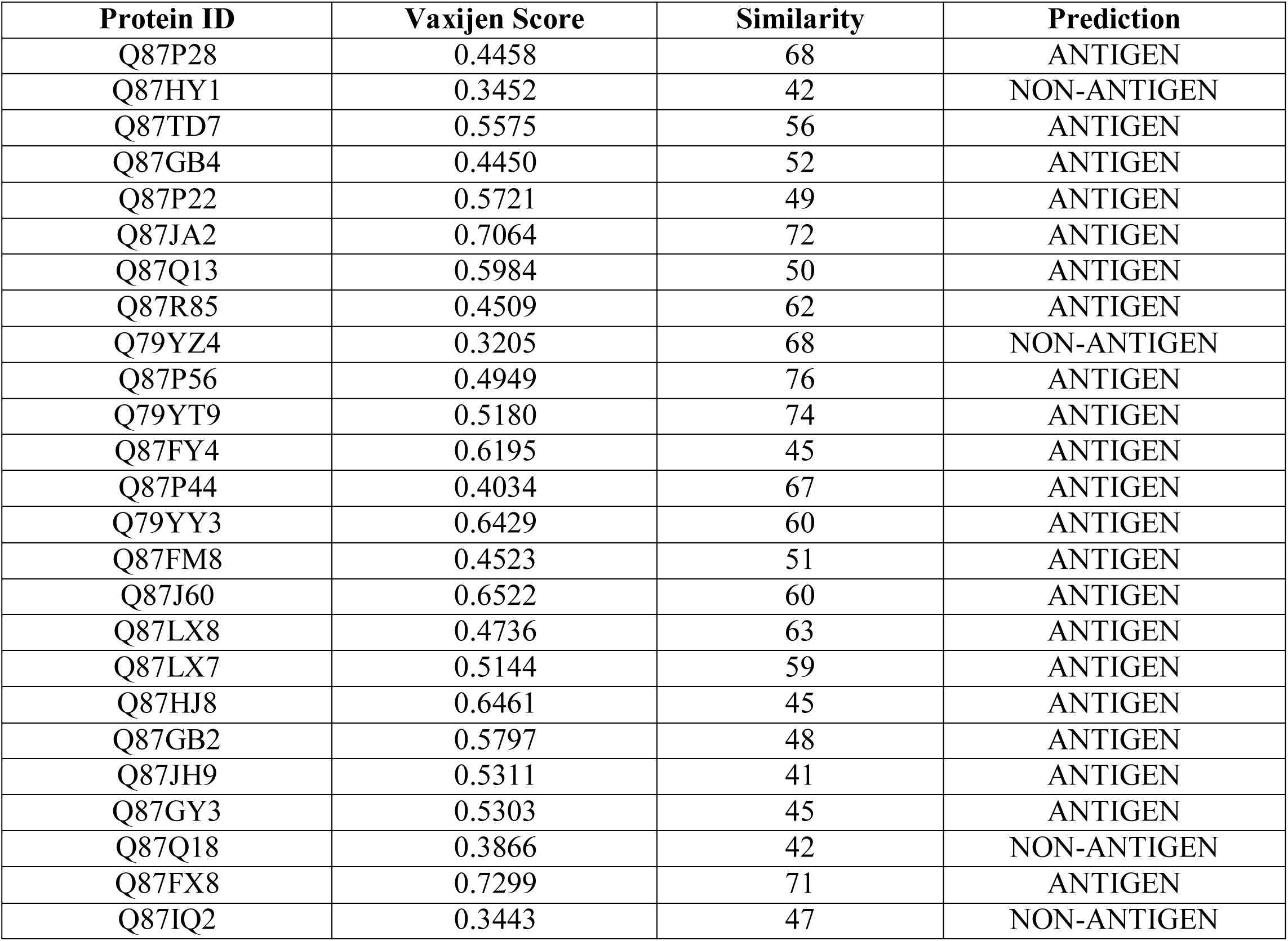
Antigenicity of novel membrane proteins (Vaccine targets) and similarity analysis with human microbiome

**Supplementary table 4:** Antigenicity and similiarity analysis of novel outer membrane proteins with human microbiome (%)

| Accession No. | Vaxijen Score | Similiarity analysis with human microbiome (%) |
| --- | --- | --- |
| Q87P28 | 0.4458 | <68 |
| Q87HY1 | 0.3452 | <42 |
| Q87TD7 | 0.5575 | <56 |
| Q87GB4 | 0.4450 | <52 |
| Q87P22 | 0.5721 | <49 |
| Q87JA2 | 0.7064 | <72 |
| Q87Q13 | 0.5984 | <50 |
| Q87R85 | 0.4509 | <62 |
| Q79YZ4 | 0.3205 | <68 |
| Q87P56 | 0.4949 | <76 |
| Q79YT9 | 0.5180 | <74 |
| Q87FY4 | 0.6195 | <45 |
| Q87P44 | 0.4034 | <67 |
| Q79YY3 | 0.6429 | <60 |
| Q87FM8 | 0.4523 | <51 |
| Q87J60 | 0.6522 | <60 |
| Q87LX8 | 0.4736 | <63 |
| Q87LX7 | 0.5144 | <59 |
| Q87HJ8 | 0.6461 | <45 |
| Q87GB2 | 0.5797 | <48 |
| Q87JH9 | 0.5311 | <41 |
| Q87GY3 | 0.5303 | <45 |
| Q87Q18 | 0.3866 | <42 |
| Q87FX8 | 0.7299 | <71 |
| Q87IQ2 | 0.3443 | <47 |

**Supplementary table 5:**
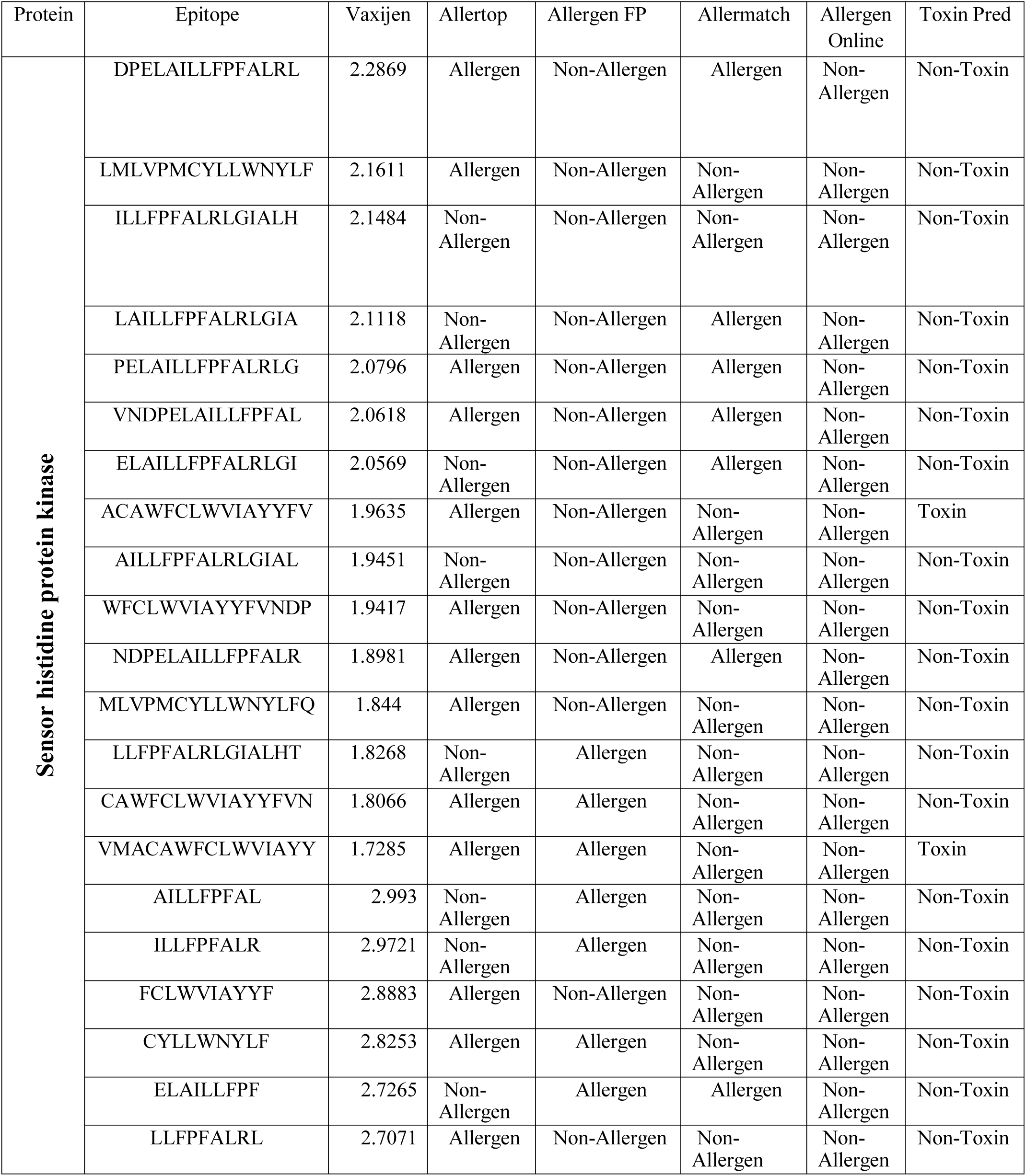
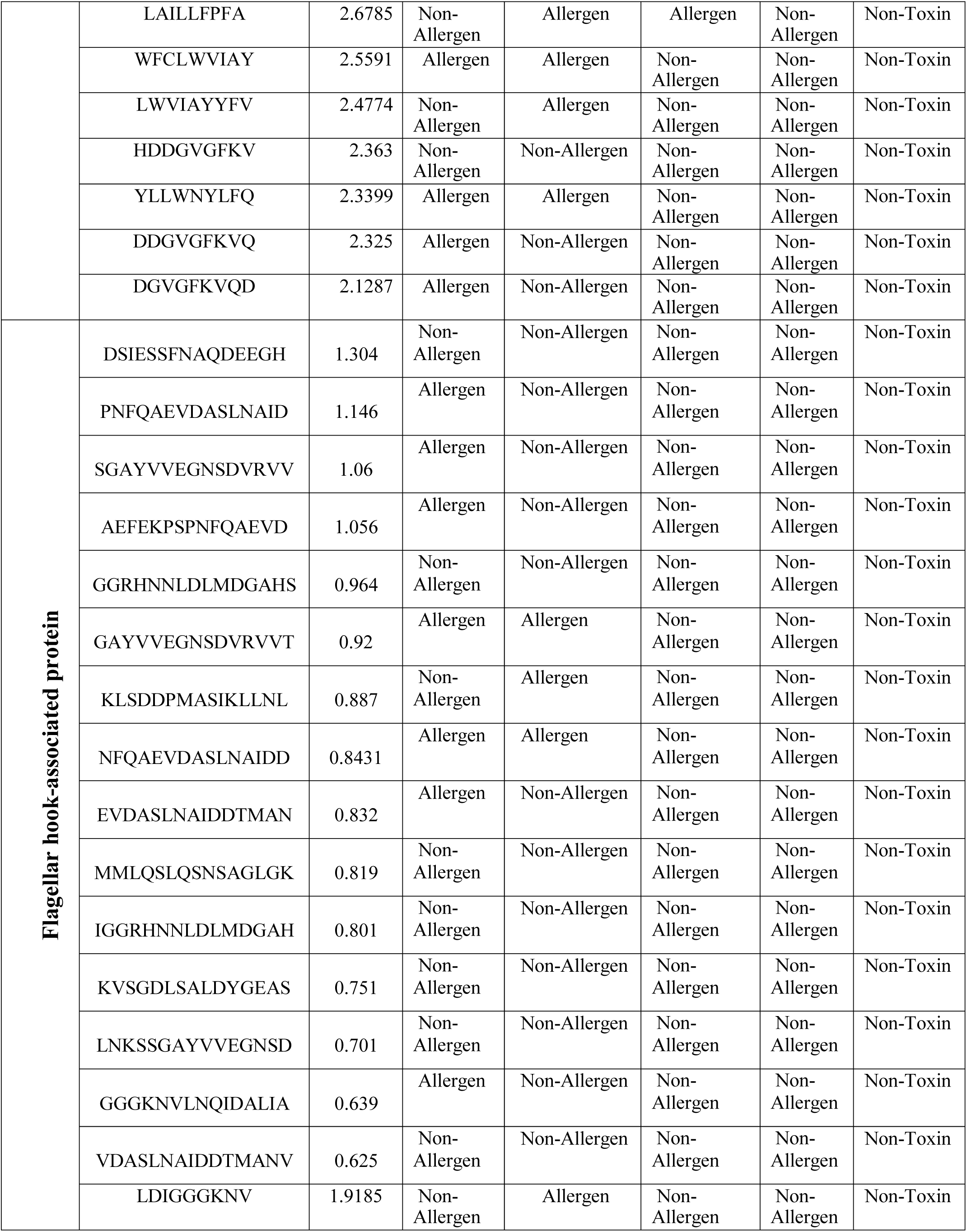
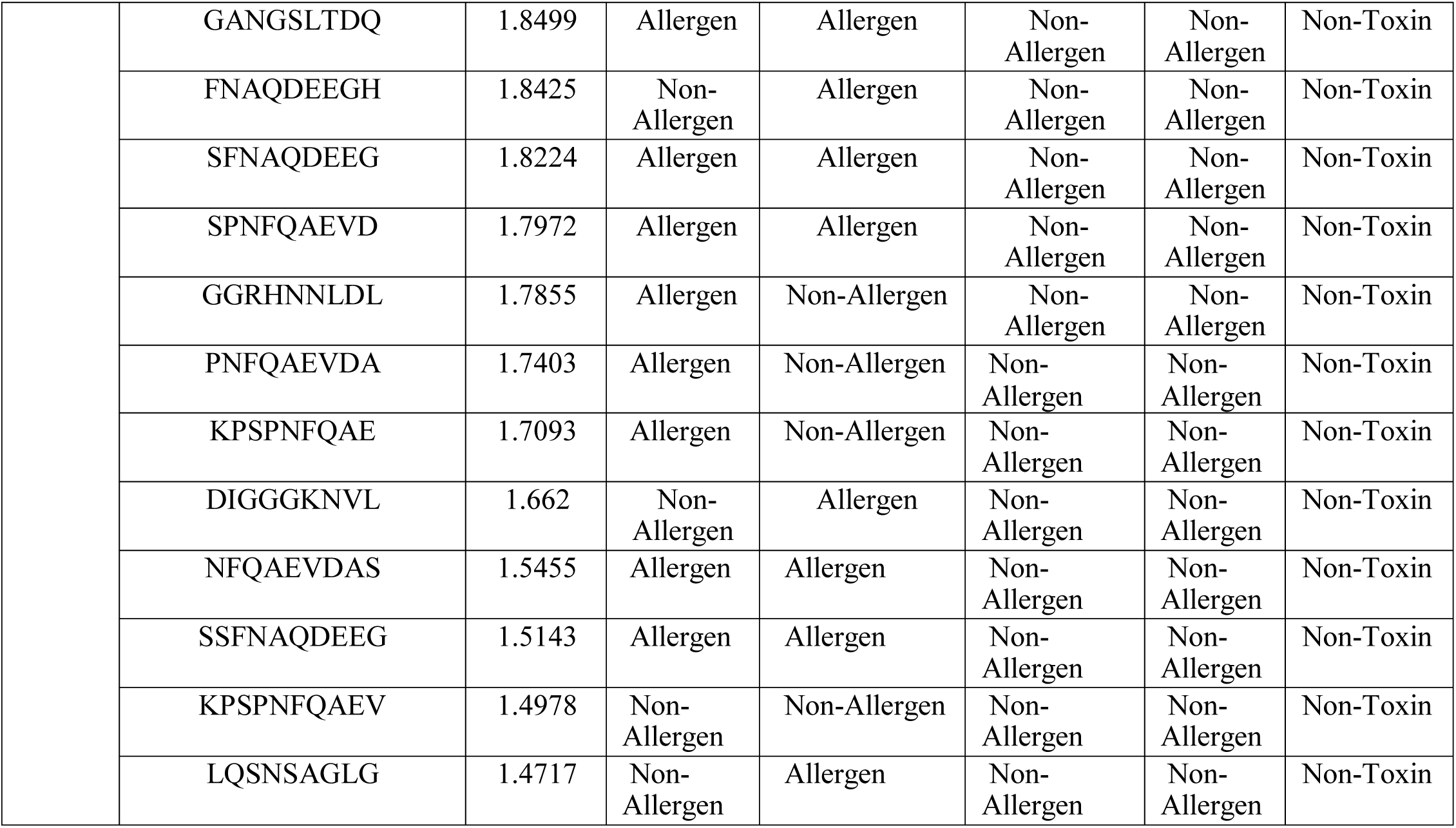
Allergenicity pattern and toxicity analysis of top epitopes for Sensor histidine protein kinase and Flagellar hook-associated protein

**Supplementary Table 6:**
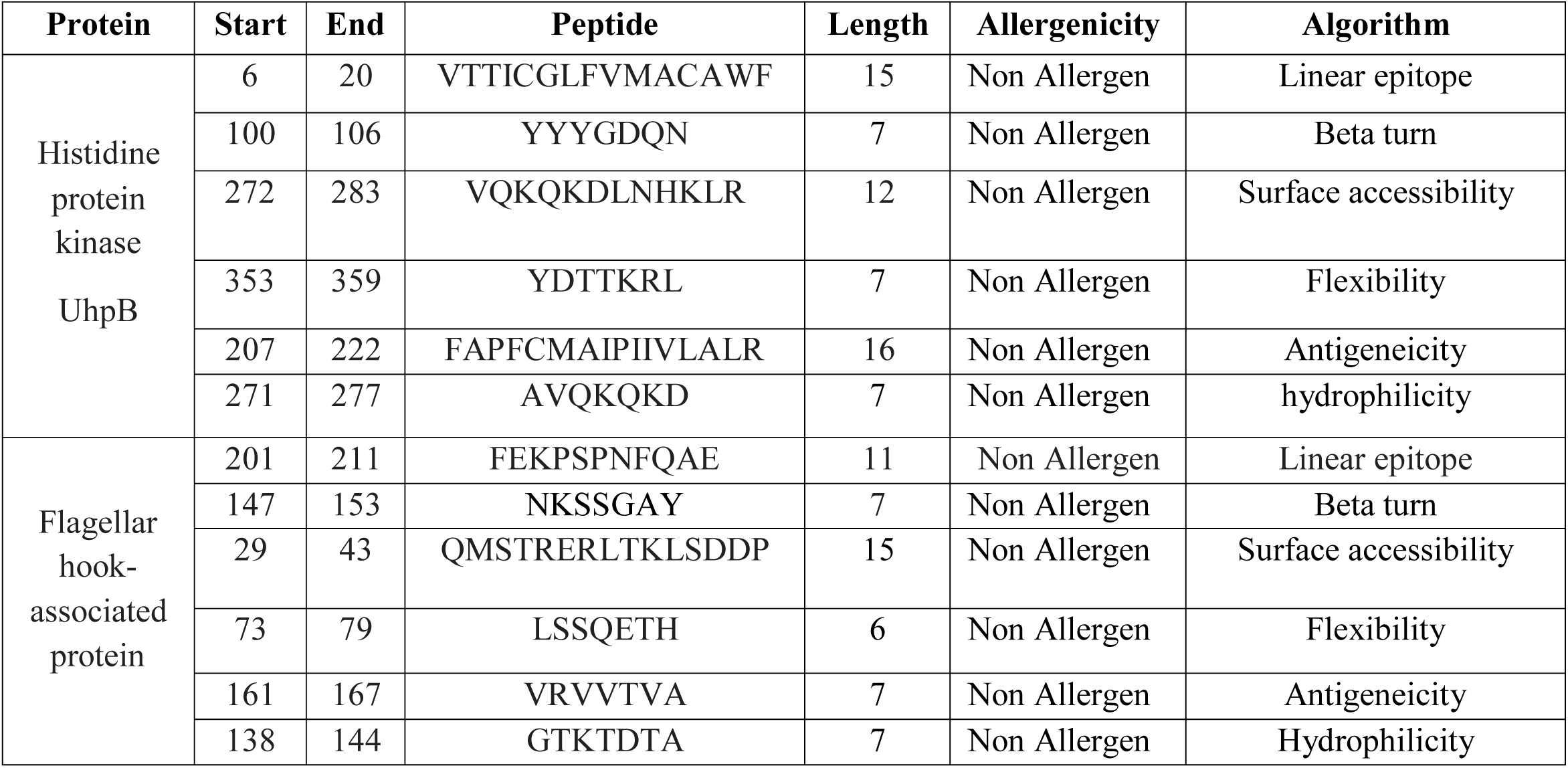
Allergenicity assessment of the predicted B-cell epitopes generated from histidine protein kinase and flagellar hook-associated protein

**Supplementary Table 7:**
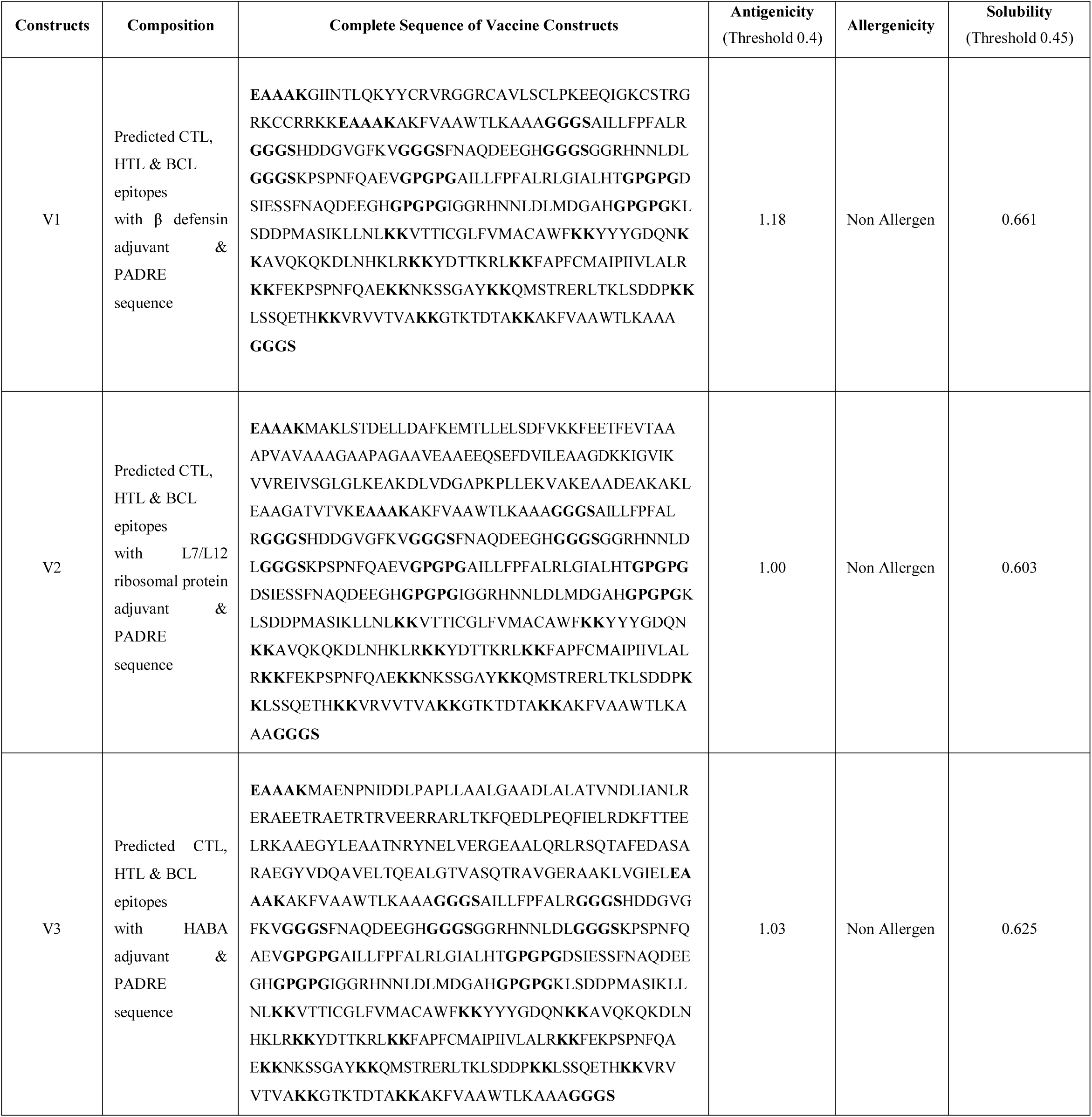
Allergenicity, antigenicity and solubility prediction of the constructed vaccines

**Supplementary Table 8:** Docking score of vaccine construct V1, V2 and V3 with different HLA alleles including HLADRB1*03:01 (1A6A), HLA-DRB5*01:01 (1H15), HLA-DRB1*04:01 (2SEB), HLA-DRB3*01:01 (2Q6W), HLA-DRB1*01:01 (2FSE) and HLA-DRB3*02:02 (3C5J).

| Vaccine Construct | HLA alleles PDB ID's | Global Energy | Hydrogen Bond Energy | ACE | Score | Area |
| --- | --- | --- | --- | --- | --- | --- |
| V1 | 1A6A | -5.77 | 0.00 | -2.93 | 15800 | 2614.80 |
|  | 1H15 | -32.57 | -1.48 | -3.79 | 16644 | 2074.50 |
|  | 2SEB | -26.95 | -2.39 | 2.40 | 17546 | 2082.60 |
|  | 2Q6W | -16.57 | -1.50 | 3.50 | 15770 | 2538.40 |
|  | 2FSE | -19.27 | -5.69 | 8.65 | 16988 | 2105.90 |
|  | 3C5J | 11.34 | -1.87 | 2.10 | 16094 | 2066.20 |
| V2 | 1A6A | -22.98 | -2.20 | -3.85 | 15152 | 2094.40 |
|  | 1H15 | -18.21 | -2.99 | 11.23 | 16964 | 2574.80 |
|  | 2Q6W | -1.13 | 0.00 | 4.11 | 18528 | 3097.60 |
|  | 2SEB | 4.86 | 0.00 | 4.62 | 17698 | 2379.80 |
|  | 2FSE | -43.58 | -5.18 | 2.59 | 16638 | 2311.70 |
|  | 3C5J | -4.64 | -1.04 | 1.32 | 18134 | 2927.80 |
| V3 | 1A6A | -0.58 | -1.24 | 4.86 | 17262 | 2533.10 |
|  | 1H15 | 11.12 | -1.51 | 8.43 | 18698 | 2572.80 |
|  | 2SEB | 17.22 | 0.00 | 3.91 | 21978 | 3151.80 |
|  | 2Q6W | -16.11 | -4.54 | 13.23 | 16068 | 2515.50 |
|  | 2FSE | -11.82 | -6.92 | 18.45 | 16308 | 2064.50 |
|  | 3C5J | -5.73 | -0.98 | 6.61 | 17746 | 2873.70 |

## Notes

### Competing Interest Statement

The authors have declared no competing interest.

### Summary of Updates

Title changed. This manuscript was submitted to PloS Computational Biology with new title

